# Different forms of variability could explain a difference between human and rat decision making

**DOI:** 10.1101/2020.01.05.895268

**Authors:** Quynh Nhu Nguyen, Pamela Reinagel

**Affiliations:** Section of Neurobiology, Division of Biological Sciences, University of California San Diego, La Jolla, California, United States of America

## Abstract

When observers make rapid, difficult perceptual decisions, their response time is highly variable from trial to trial. In a visual motion discrimination task, it has been reported that human accuracy declines with increasing response time, whereas rat accuracy increases with response time. This is of interest because different mathematical theories of decision-making differ in their predictions regarding the correlation of accuracy with response time. On the premise that perceptual decision-making mechanisms are likely to be conserved among mammals, we seek to unify the rodent and primate results in a common theoretical framework. We show that a bounded drift diffusion model (DDM) can explain both effects with variable parameters: trial-to-trial variability in the starting point of the diffusion process produces the pattern typically observed in rats, whereas variability in the drift rate produces the pattern typically observed in humans. We further show that the same effects can be produced by deterministic biases, even in the absence of parameter stochasticity or parameter change within a trial.

## Introduction

One might expect decision-making by humans to be quite different from that of rats. In decisions with wide-reaching long-term consequences, we expect (or at least wish) humans would avail themselves of abstract conceptual thought, logical reasoning, and culturally accumulated knowledge that would be unavailable to a rat. Yet all organisms face a continuous challenge of selecting among alternative available actions in order to pursue goals. In order to select an action, sensory information, internal knowledge, and goals are combined to assess and evaluate the likely outcomes of possible actions relative to survival needs. Often there is not enough time to acquire the evidence necessary to determine with certainty the optimal course of action, so an action must be selected despite unresolved or unresolvable uncertainty. Some mechanism is needed to ensure timely commitment and to optimize outcome on average, and this must adapt flexibly to prevailing sensory context, shifting goal priorities, the urgency of action, and the severity of consequences of errors. When it comes to the continuous sensory guidance of moment-by-moment actions, decisions about sensory evidence are made in a fraction of a second. We speculate that in this case, mechanisms are largely conserved across mammals.

A now-classic series of studies in humans and non-human primates introduced the use of a stochastic visual motion task to study decision making (Britten, Shadlen et al. 1992, Britten, Shadlen et al. 1993, Britten, Newsome et al. 1996, Shadlen, Britten et al. 1996, Shadlen and Newsome 1996, Gold and Shadlen 2001, Shadlen and Newsome 2001, Roitman and Shadlen 2002, Mazurek, Roitman et al. 2003, Huk and Shadlen 2005, Palmer, Huk et al. 2005, Gold and Shadlen 2007). In each trial a visual stimulus provides information regarding which of two available actions is associated with reward and which is associated with non-reward or penalty. Stimulus strength is modulated by the motion coherence, which is defined as the fraction of the dots in the display that are “signal” (moving toward the rewarded response side). The remaining dots are “noise” (moving in random directions). As stimulus strength increases, accuracy increases and response time decreases for both monkeys (Roitman and Shadlen 2002) and humans (Palmer, Huk et al. 2005). This is parsimoniously explained by drift diffusion models, which postulate that noisy sensory evidence is integrated over time until the accumulated evidence reaches a decision threshold (Stone 1960, Ashby 1983, Busemeyer and Townsend 1993, Gold and Shadlen 2001, Usher and McClelland 2001, Ratcliff and Tuerlinckx 2002, Palmer, Huk et al. 2005, Gold and Shadlen 2007, Brown and Heathcote 2008, Ratcliff and McKoon 2008, Ratcliff, Smith et al. 2016). Although this class of model is highly successful, more data are needed to test model predictions and differentiate among competing versions of the model and alternative model classes (Wang 2002, Ratcliff and McKoon 2008, Pleskac and Busemeyer 2010, Purcell, Heitz et al. 2010, Rao 2010, Heathcote and Love 2012, Tsetsos, Gao et al. 2012, Huang and Rao 2013, Usher, Tsetsos et al. 2013, Scott, Constantinople et al. 2015, Ratcliff, Smith et al. 2016, Sun and Landy 2016, White, Servant et al. 2018).

For example, when monkeys or humans perform this task, among trials of the same stimulus strength the interleaved trials with longer response times are more likely to be errors (Roitman and Shadlen 2002, Palmer, Huk et al. 2005). In its simplest form the drift diffusion model does not explain this result; therefore the observation has been an important constraint for recent theoretical efforts. The result can be explained if the decision bound is not constant but instead decays as a function of time (Churchland, Kiani et al. 2008, Cisek, Puskas et al. 2009, Bowman, Kording et al. 2012, Drugowitsch, Moreno-Bote et al. 2012). A collapsing decision bound can be rationalized as an optimal strategy under some task constraints (Rao 2010, Hanks, Mazurek et al. 2011, Huang and Rao 2013, Tajima, Drugowitsch et al. 2016) though this argument has been challenged by others (Hawkins, Forstmann et al. 2015, Boehm, Hawkins et al. 2016). There are alternative ways to explain the data within the sequential sampling model framework without positing an explicit urgency signal or decaying bound (Ditterich 2006, Ditterich 2006, Ratcliff and McKoon 2008, Ratcliff and Starns 2013).

When rats performed the same random dot motion task, however, the opposite effect was found: their later decisions were more likely to be accurate (Reinagel 2013, Shevinsky and Reinagel 2019). The same has also been reported for image discriminations in rats (Reinagel 2013), for visual orientation decisions in mice (Sriram, Li et al. 2020), and in humans in some other tasks (McCormack and Swenson 1972, Ratcliff and Rouder 1998, Long, Jiang et al. 2015, Stirman, Townsend et al. 2016). This result is not readily explained by some of the models suggested to explain the late errors of primates (reviewed in (Heitz 2014, Ratcliff, Smith et al. 2016, Hanks and Summerfield 2017)). Here we explore a stochastic variant of the drift-diffusion model (Ratcliff and Tuerlinckx 2002, Ratcliff and McKoon 2008) for its ability to explain these problematic findings in both species.

## Results

In a basic drift diffusion model (DDM), the relative sensory evidence in favor of a decision (e.g. “motion is rightward” vs. “motion is leftward”) is accumulated by an internal decision variable, resulting in a biased random walk, i.e., diffusion with drift (Fig 1A). The average drift rate is determined by the sensory signal strength (e.g., the coherence of visual motion). When the decision variable reaches either decision threshold, the agent commits to a choice. The time at which the decision variable crosses a threshold (response time), and the identity of the decision threshold that is crossed (correct vs. incorrect), vary from trial to trial. The model parameters are the starting point *z*, threshold separation *a*, drift rate *v*, and nondecision time *t* (in Fig 1A-E, *z* = 0 *a* = 2, *t* = 0, *v* = 0.7).

**Figure 1.**
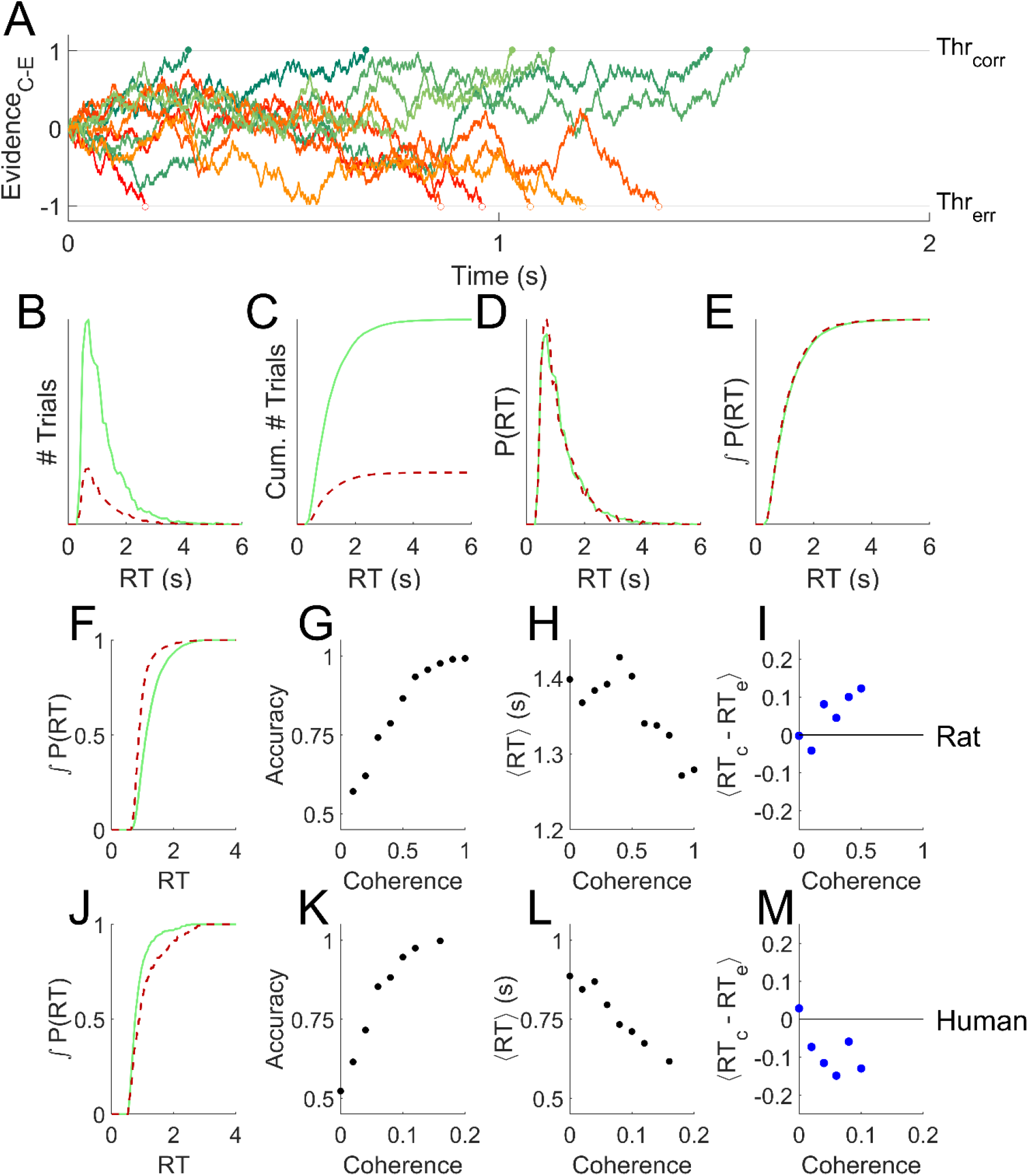
The basic drift diffusion model is incompatible with data from rats or humans. **(A)** Simulated evidence accumulation in a basic drift diffusion model, for six example trials that terminated in correct decisions (green shades, solid symbols) and six that terminated in error decisions (red shades, open symbols). Although an equal number of traces are shown, in the condition illustrated 80% of the traces terminated by crossing the correct boundary. **(B)** Response time distributions for errors (red, dashed) vs. correct trials (green, solid) of 10^4^ trials like those simulated as in A. **(C)** Cumulatives of the distributions shown in B. **(D-E)** Distributions in B and C normalized to the number of trials. **(F)** Cumulative probability distribution of response time for errors and correct trials for an example experiment in a rat, a fixed-coherence experiment with 65% coherence, for which the rat was 82% correct. The null hypothesis that the distributions are the same can be rejected with P=2.91e-171 (N=8851,1913) by two-tailed Kolmogorov–Smirnov (KS) test. **G-I.** Analysis of an example rat psychometric experiment. **(G)** Accuracy increased with coherence. **(H)** Mean response time decreased with coherence. **(I)** On average, the response time of a correct trial is greater than that of a temporally nearby error trial of the same coherence. **(J)** Like F, for an example experiment in a human, a fixed-coherence experiment with 10%coherence for which the subject was 83% correct; P= 4.69e-05 (N=745,153) by KS test. Errors are later than correct trials, unlike either the DDM model (E) or rats (F). **(K-M)** Like G-I, for an example human psychometric experiment. **(M)** Error trials are longer than correct trials on average, unlike rats, and also incompatible with DDM. Data replotted from (Shevinsky and Reinagel 2019, Reinagel and Shevinsky 2020).

An interesting feature of this model is that for any set of parameters, the errors and correct responses have identical response time distributions (Fig 1B-E, red vs. green). Therefore errors are on average the same speed as correct responses – even if the signal is so strong that errors are very rare.

We note that this does not, but may at first appear to, contradict two other facts. First, responses to stronger stimuli tend to be both more accurate and faster, which in this model is explained by a higher drift rate *v*. In this sense response time is negatively correlated with accuracy – but only when comparing trials of differing stimulus strengths. Second, conservative subjects tend to take more time to respond and are more accurate, which in this model is explained by a greater threshold separation *a*. In this sense response time is positively correlated with accuracy – but only when comparing blocks of trials with different overall degrees of caution. Both of these facts are consistent with the fact that within a block of fixed overall caution, comparing among the trials of the same stimulus strength, response time and accuracy are uncorrelated in the basic DDM model.

Both humans and rats deviate systematically from the prediction that correct and error trials have the same mean and probability distribution, however (Shevinsky and Reinagel 2019). In the random dot motion discrimination task, for example, correct trials of rat subjects tend to have longer response times compared to errors (e.g., Fig 1F, cf. 1E). We quantify this effect by comparing the response times of individual correct trials to nearby (but not adjacent) error trials of the same stimulus strength (Fig 1I). This temporally local measure is robust to data non-stationarities that could otherwise produce a result like that shown in Fig 1F artefactually (Shevinsky and Reinagel 2019). Humans also violate the basic DDM model prediction, but in the opposite way. For humans, errors tend to have longer response times (e.g., Fig 1J, cf.1E; summarized in 1M). Our goal is to find a unified framework to account for both these deviations from predictions.

### Drift Diffusion Model with variable parameters

It was previously shown that adding noise to the parameters of a bounded drift diffusion model can differentially affect the error and correct response time distributions (Ratcliff and Tuerlinckx 2002, Ratcliff and McKoon 2008). The version we implemented has three additional parameters: variability in starting point σ_z_, variability in non-decision-time σ_t_, and variability in drift rate σ_v_ (Fig 2A). We are able to find parameter sets that produce behavior qualitatively similar to either a rat (Fig 2B-E, cf. Fig 1F-I) or a human (Fig2 F-I, cf. Fig 1J-M). Notably, this model can replicate the shift in the response time distribution of correct trials to either later or earlier than that of error trials (Fig 2B, cf. Fig 1F; Fig 2F, cf. Fig2J), unlike the standard DDM (Fig 1E). The model also replicates the fact that the amplitude of this effect increases with stimulus strength (Fig 2E solid blue symbols, cf. Fig 1I; Fig 2I, cf. Fig 1M). Removing the drift rate variability and starting point variability from these simulations improved accuracy (Fig 2C,G open symbols), increased the response time for ambiguous stimuli (Fig 2D,H), and eliminated the average difference between correct and error response times (Fig 2E,I).

**Figure 2.**
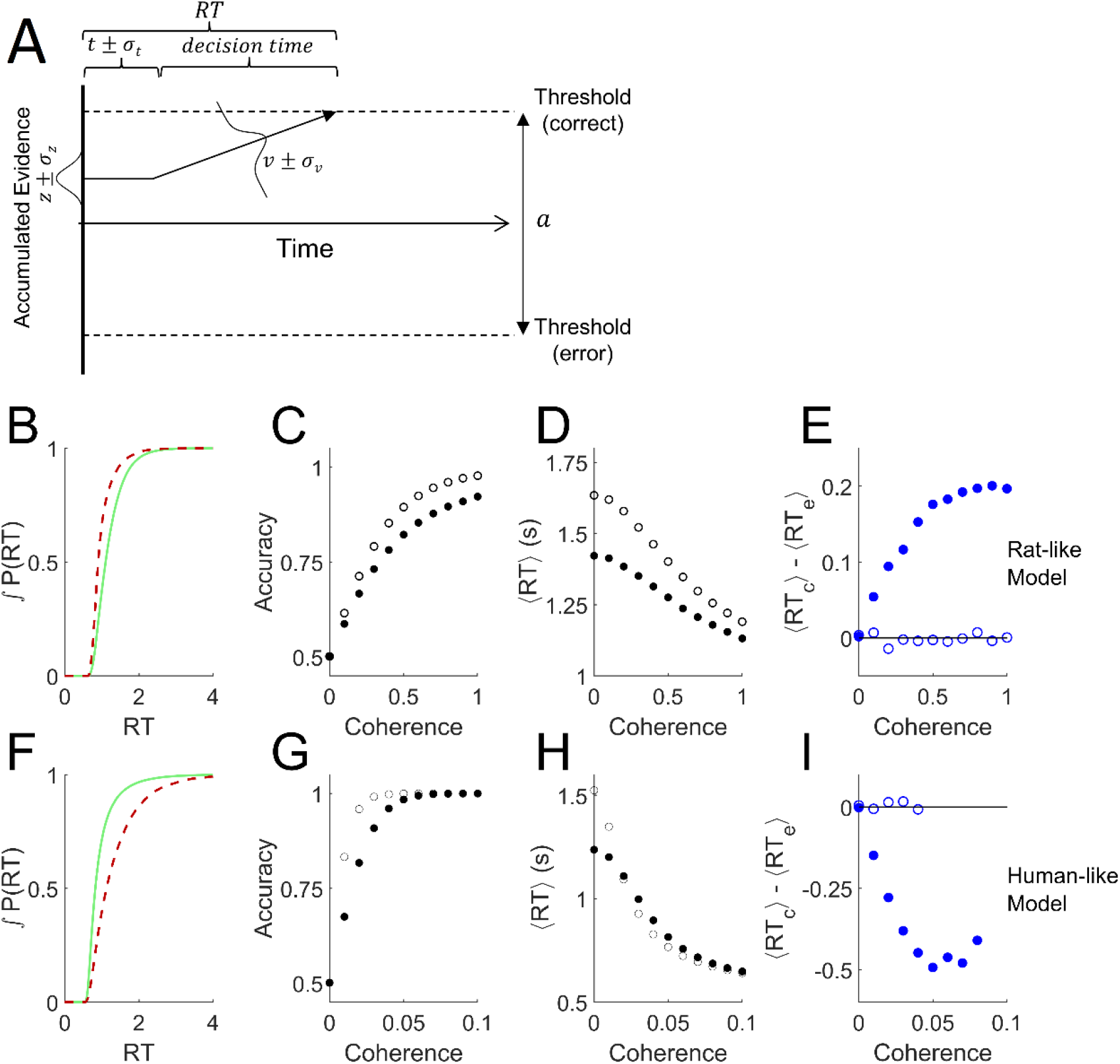
Addition of variability to the parameters of the drift diffusion model. **(A)** Definition of parameters. The parameter *a* is the distance between the error and correct thresholds. The starting point of evidence accumulation is given by *z*. The average drift rate *v* depends on the stimulus strength. The observable outcomes are response time (RT) and decision (correct or error). The non-decision-time *t* reflects both sensory latency and motor latency, but is drawn at left for graphical simplicity. Parameters *t*, *z*, and *v* vary from trial to trial according to the variability parameters σ_t_, σ_z, and_ σ_v_ respectively. Drift rate variability was simulated by a normal distribution around the mean parameter. Starting point variability and non-decision time variability were simulated by uniform distributions centered on the mean parameter. Diagram after (Ratcliff and McKoon, 2008). **(B-E)** Analysis of trials simulated with parameters that produce qualitatively rat-like behavior: *a* = 1.84, *t* = 0.74, *σ_v_* = 0.1, *σ_z_* = 1.5, *σ_t_* = 0.2, and *v* = −0.5 *c*^2^ + 2.5*c*, where *c* is coherence (motion stimulus strength). **(B)** Cumulative probability distributions of correct vs. error trial response times, for *c* = 0.8. **(C)** Psychometric curve. **(D)** Chronometric curve **(E)** Difference between mean response times of errors and correct trials. **(F-I)** Like panels B-E but with parameters that produce qualitatively human-like behavior: *a* = 2.0, *t* = 0.5, *σ_v_* = 1.5, *σ_z_* = 0.3, *σ_t_* = 0.03, and *v* = −80 *c*^2^ + 80*c*. **(F)** Cumulative RT probability distributions for *c* = 0.03. For all these simulations, the mean starting point *z* = 0, timestep *τ* = 0.001, diffusion noise *σ_n_* = 1, *N* = 10^5^ trials per coherence. Open symbols show results obtained after setting σ_z_=0 and σ_v_=0. Note that in conditions with 100% accuracy the 〈*RT_c_*〉 − 〈*RT_e_*〉 difference is undefined.

We systematically varied the parameters of this model (Fig 3A-C) to determine all the conditions under which the mean RT of correct trials can be greater or less than the mean RT of error trials, using parameter ranges from the literature (Ratcliff and Tuerlinckx 2002, Wagenmakers, van der Maas et al. 2007, Ratcliff and McKoon 2008). Like the basic DDM, the simulations with *σ_z_* = 0, *σ_v_* = 0 showed no difference between correct and error RT for any drift rate (black curves are on y=0 line), in spite of the addition of non-decision time variability *σ_t_*. We never observed a positive RT difference in this model unless the starting point was variable (the dark blue or black curves, *σ_z_* = 0, lie entirely on or below the abscissa). Whenever *σ_z_* > 0 and *σ_v_*=0, the RT difference was positive (thin lines other than black). We never observed a negative RT difference in the absence of drift rate variability (thin lines, *σ_v_* = 0, lie entirely on or above the abscissa). Whenever *σ_v_* > 0 and *σ_z_*=0, the RT difference was negative (blue curves). Holding other parameters constant, the RT difference always increased (more positive, or less negative) with increasing *σ_z_* (blue→red) and decreased with increasing *σ_v_* (thin → thick).

**Figure 3.**
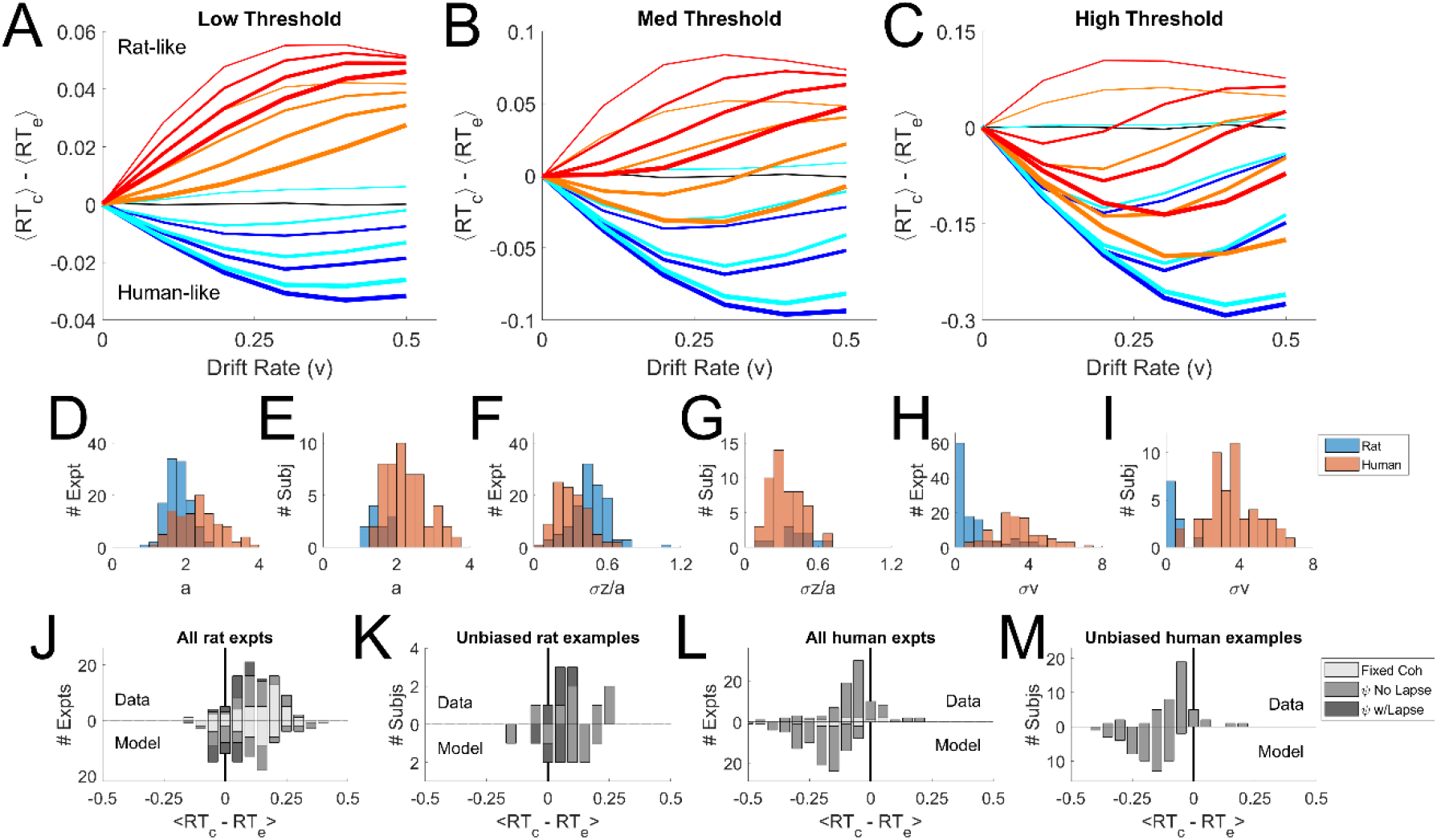
Parameter sweep of the variable-parameter DDM. **(A-C)** Curves show the difference between correct and error mean response times, 〈*RT_correct_*〉 − 〈*RT_error_*〉, as a function of drift rate parameter *v* = [0.0,0.1,0.2,0.3,0.4,0.5]. Colors indicate starting point variability *σ_z_* = [0.0,0.02,0.07,0.10], in spectral order from low (dark blue) to high (red). Line thickness indicates drift rate variability *σ_v_* = [0.0,0.08,0.12,0.16]. Black curve is for *σ_v_* = 0, *σ_z_* = 0. **(A)** Simulations with threshold separation *a* = 0.08. **(B)** Simulations with *a* = 0.11. **(C)** Simulations with *a* = 0.16. For all the simulations shown, *z* = 0, *t* = 0.3, *σ_t_* = 0.2, *τ* = 0.001, and *σ_n_* = 0.1. **(D)** Distribution of threshold separation *a* from fits of this model to datasets from (Reinagel and Shevinsky 2020). **(E)** Distribution of *a* among example psychometric experiments from unique unbiased subjects in those datasets (see Methods). **(F)** Distribution of starting point variability, expressed as a fraction of threshold separation: *σ_z_/a*. **(G)** Distribution of *σ_z_/a* in the unbiased example sets. **(H)** Distribution of drift rate variability *σ_v_*. **(I)** Distribution of *σ_v_* in the unbiased example sets. **(J)** Average difference between correct and error response times 〈*RT_c_* − *RT_e_*〉 computed locally within coherence, averaged over coherences ≥0.4. Fixed coherence (light), psychometric with <10% lapse (medium) or with ≥10% lapse (dark). Upward bars show the results from the rat dataset, preplotted from (Shevinsky and Reinagel 2019); N=51 psychometric, N=38 fixed-coherence experiments had sufficient trials for this analysis. Lower bars show results for trial data simulated by the models (N=58 psychometric and N=39 fixed). **(K)** Like J, but for the rat unbiased example subset; N=11 data or models. **(L)** Like J but for human dataset (N=81 psychometric, N=9 fixed) and models fit to human dataset (N=93 psychometric,N=9 fixed), averaged over coherences ≥0.04. **(M)** Like L but for the human unbiased example subset, N=45 (data) or N=51 (model). For descriptive statistics see Supplemental Information. Because of the limitations of fitting, we refrain from making statistical claims about comparisons of human-to-rat or data-to-model distributions.

When both starting point variability and drift rate variability are present simultaneously, these opposing effects trade off against one another quantitatively, such that there are many parameter combinations consistent with any given sign and amplitude of effect. This explains why parameter fits to data are generally degenerate.Taken together, the simulations show that human-like pattern is associated with dominance of *σ_v_* and the rat-like pattern with dominance of *σ_z_*. The nondecision time *t* and its variability *σ_t_* were explored in separate simulations and did not impact the effect of interest (not shown).

It is difficult or impossible to recover the true parameters of this model by fitting data (Boehm, Annis et al. 2018). Nevertheless, we fit published human and datasets to identify example parameters of the model consistent with the observed data. The parameters obtained from fitting are not guaranteed to be the optimal solutions of the model nor accurate measures of noise in the subjects. Bearing these caveats in mind, the distributions of parameters we obtained (Fig 3 D,F, and H) were consistent with the parameter sweeps. Parameters fit to humans and rats overlapped substantially, but the human distributions were shifted toward those that favor late errors (higher *σ_v_*, higher *a* and lower *σ_z_*), and rats’ parameters toward those that favor early errors (higher *σ_z_*, lower *a* and *σ_v_*). Trials simulated using the fitted parameters reproduced the sign of the effect (Fig 3J,L). In a subset of examples defined by low bias, the difference between parameter distributions of rats and humans were less pronounced (Fig 3 E,G, and I). Experiments with low bias still exhibit the species difference, and models fit to those examples still reproduced the species difference (Fig 3K,M).

This analysis does not prove that humans and rats have trial-by-trial variability in drift rate and starting point, much less provide an empirical measure of that variability. What it does show is that if starting point and drift rate vary from trial to trial, that alone could be sufficient to produce the effects previously reported in either species. Only subtle differences in the relative dominance of drift rate vs. starting point variability would be required to explain the reported species difference.

### Variability need not be random

Random trial-to-trial variability in parameters can cause differences between correct and error response times. But “variability” does not have to be noise. Systematic biases in the starting point or drift rate would also vary from trial to trial, and therefore would produce similar effects. We tested whether bias alone could produce results resembling those of (Shevinsky and Reinagel 2019).

First, recall that we have defined decision thresholds as “correct” vs. “error” rather than “left” vs. “right” (Fig 2A). Therefore it is impossible for the mean starting point *z* to be biased, because the agent cannot know *a priori* which side is correct. If a subject’s starting point were systematically biased to one response side, the starting point would be closer to the correct threshold on half the trials (when the preferred side was the correct response), but further on the other half of the trials (when the non-preferred response was required). Thus the mean starting point would be *z* = 0, but the distribution of *z* would be binary (or at least bimodal), and thus high in variance. This could mimic a model with high *σ_z_*, even in the absence of stochastic parameter variability.

We demonstrate by simulation that adding a fixed response-side bias to the starting point of an otherwise standard DDM is sufficient to produce a response bias (Fig 4A). Response accuracy is higher when the correct response (“target”) is on the preferred side, and can fall below chance for weak stimuli to the non-preferred side (Fig 4B). At any given coherence, response times are faster for targets on the preferred side (Fig 4C). For targets on the preferred side, reaction times of correct trials are faster than errors, whereas for targets on the nonpreferred side, correct trials are slower than errors (Fig 4D). This is because when the target is on the preferred side, the starting point is closer to the correct threshold, such that correct responses cross threshold faster than error responses, while the opposite is true for targets on the nonpreferred side. Thus both the left and right target trials violate the expectation of 〈*RT_c_*〉 = 〈*RT_e_*〉, but in opposite directions.

**Figure 4.**
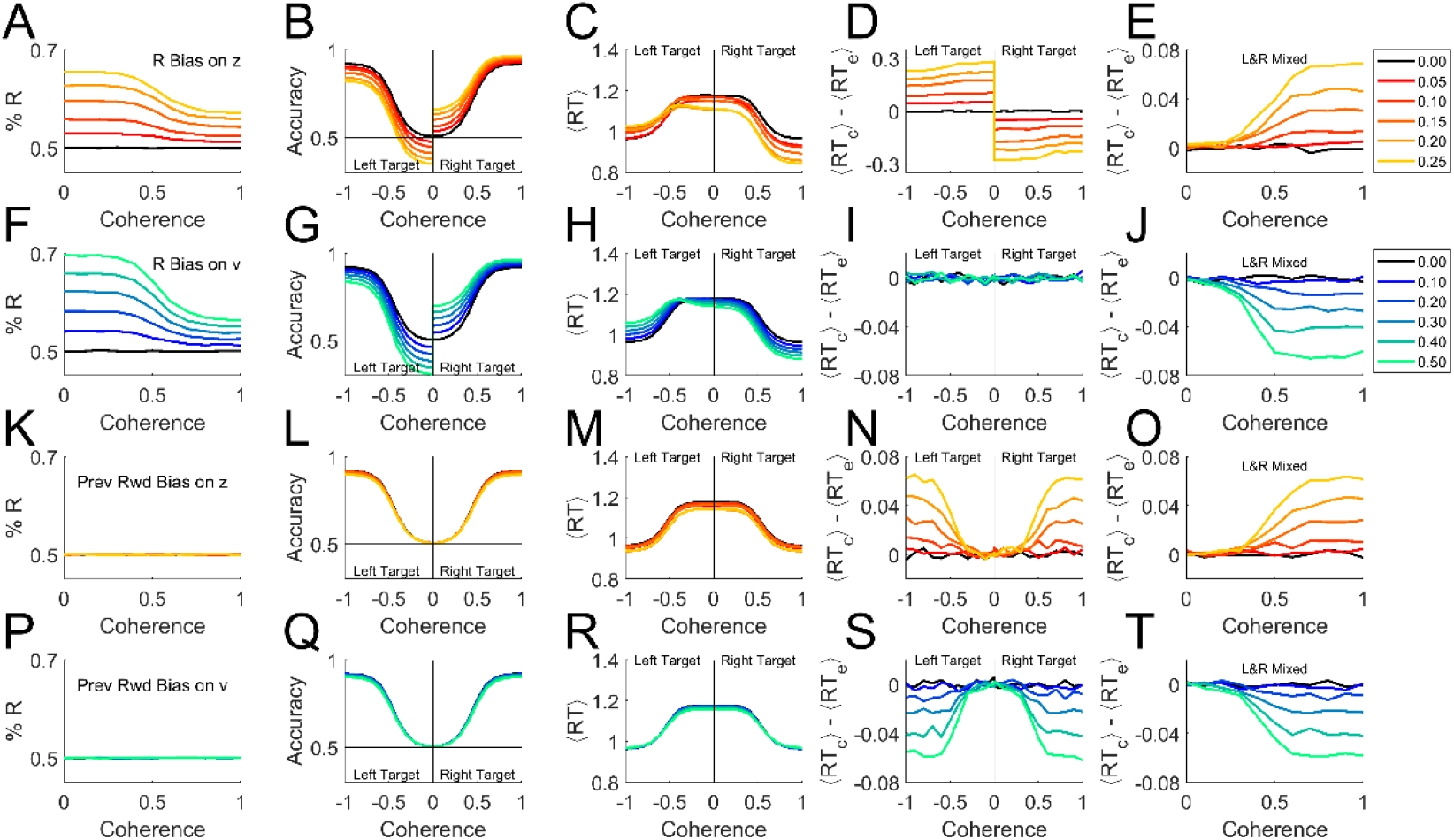
Bias is sufficient to produce either a rat-like or human-like effect within the basic DDM. Simulations were performed with *σ_v_* = 0, *σ_z_* = 0, *σ_t_* = 0 (i.e. the basic model in Fig 1) but with different forms of bias added. Trials were simulated with 50% rightward motion (right response targets), with 5×10^5^ trials per coherence. **(A-E)** A right side bias was simulated by displacing the starting point *z* toward the correct boundary on right-target trials, or toward the error boundary for left-target trials, by the amount indicated by color key at right. Where coherence is signed, negative indicates leftward motion and positive, rightward motion. If a coherence axis is unsigned, the left and right motion trials are pooled. **(A)** Percent right responses, as a function of coherence (sensory stimulus strength), which determines the drift rate *v*. **(B)** Average accuracy of the response as a function of coherence. **(C)** Average response time as a function of coherence. **(D)** Difference between correct and error mean response times, 〈*RT_correct_*〉 − 〈*RT_error_*〉, a function of coherence. **(E)** Difference between correct and error mean responses times when left-motion and right-motion trials are pooled. Compare to rat data (Fig 1I) or high *σ_z_* model (thin red curves in Fig 3A-C). **(F-J)** like A-E, but here bias was simulated by increasing the drift rate *v* on R-target trials, or decreasing it on L-target trials, by the amount indicated in color key at right. Compare panel J to human data (Fig1M) or high *σ_v_* model (thick blue curves in Fig 3A-C). **(K-O)** like A-E but here the starting point *z* was displaced toward the side that was rewarded in previous trial (if any). Same color key as A-E. **(P-T)** like F-J but here the drift rate *v* was increased if the target was on the side rewarded in previous trial, or decreased if the target was on the opposite side. Same color key as F-J. For a similar analysis separated by behavioral choice instead, see Supplementary Information, Fig S2.

If the left-target and right-target trials are pooled in a single analysis – even if exactly equal numbers of both kinds are used – these opposite effects do not cancel out (Fig 4E). On average, correct trials would have longer RT than error trials, to an increasing degree as coherence increases (Fig 4E), just as commonly seen in rodents (e.g. Fig 1I; see (Shevinsky and Reinagel 2019)). The reason for this, in brief, is that the side with the starting point nearer the error threshold is responsible for the vast majority of the errors, and these errors have short RTs. The imbalance of contributions to correct responses is less pronounced. Although the side with the starting point nearer the correct threshold contributes the majority of correct responses, and those have short RTs, the drift rate ensures that both sides contribute substantial numbers of correct trials. For a more detailed account see Fig S1. Mechanisms aside, the important point is that if a response bias is present (Fig 4A) and an effect like that in Fig 4E is obtained from an analysis that pools left- and right-target trials, starting point bias toward the preferred side should be considered as a possible cause. Either the analysis shown here (Fig 4D) or one that separates left-side from right-side choices (Fig S2D) can be used to reveal the contribution of starting-point bias to early errors.

What if response bias arose, not from a shift in the starting point of evidence accumulation, but rather from an assymetry in the drift rate for leftward vs. rightward motion: *v_R_* ≠ *v_L_*? Again, there would be an excess of responses to the preferred side (Fig 4F), and preferred target trials would be more accurate (Fig 4G) and faster (Fig 4H). If left and right targets were analyzed separately, each on its own would have a fixed drift rate, and therefore would behave as predicted by DDM: correct trials and errors would have the same mean reaction time (Fig 4I).

But if left- and right-target trials were pooled, *v* would be biased toward or away from the correct response in different trials with equal probability, resulting in a binary or bimodal distribution in *v*. Thus the standard deviation of *v* (*σ_v_*) would be large, producing effects equivalent to high drift rate variability *σ_v_* (Fig 4J), just as commonly seen in primates (e.g. Fig 1IM; see (Shevinsky and Reinagel 2019)). The reason for this is that pooling left- and right-target trials is equivalent to mixing together trials from high- and low-coherence stimuli: the slower RTs over-represent the low-coherence (slow, inaccurate) trials while faster RTs over-represent the high coherence (fast, accurate) trials, such that errors are on average slower than correct trials (Fig S1). Therefore, if a response bias is present (Fig 4F) and an effect like that in Fig 4J is observed in a pooled-trial analysis, drift rate bias is a candidate mechanism. Either the analysis shown here (Fig 4I) or one that separates left-side from right-side choices (Fig S2I) can be used to clarify the contribution of drift rate bias to late errors.

Finally, note that rats and humans with the same degree of response-side bias could have opposite effects on 〈*RT_correct_*〉 − 〈*RT_error_*〉 (Fig 4E vs. J), if the starting point were more biased in rats and drift rate more biased in humans.

We belabor the effects of response-side bias in order to draw a broader generalization. The results just shown (Fig 4A-J) require only that a bias to one side (L or R) exists in each trial. It does not matter if that bias is fixed or varying from trial to trial, only that it is uncorrelated with the correct response. If the starting point or drift rate were biased in individual trials based on the recent trial history, for example, this would also bias the decision towards or away from the correct response with equal probability in each trial. Therefore history-dependent bias can also mimic either high *σ_z_* (Fig 4O) or high *σ_v_* (Fig 4T). But in this case, there would be no overall left or right side bias (Fig 4K,P), and even after conditioning the analysis on the target side, the “early error” (Fig 4N) or “late error” (Fig 4S) phenotypes would persist. By analogy to the case of response side bias, one could test for this specific kind of bias by conditioning the analysis on the location of the previous trial’s reward.

In principle, therefore, biases due to trial history or other contextual states could also be sufficient to explain the observed difference between error and correct response times in both species, even in the absence of overt side bias, random variability of parameters, or within-trial parameter change. Again, the difference between rats and humans does not require that historical or contextual biases are stronger in either species, only that when present, they have a stronger effect on drift rate in humans and a stronger effect on starting point in rats.

In real data, however, the observed effects could be explained by a combination of response-side bias, history-dependent bias, contextual modulation, and noise, impacting both starting point and drift rate in both species. Therefore, conditioning the analysis on discrete trial types is not a practical way to detect (or rule out) bias effects in most data sets. Other new modeling approaches show promise for dissecting such mixed effects, however (Urai, de Gee et al. 2019, Ashwood, Roy et al. 2020).

## Discussion

The impetus for this study was an observed difference between primate and rodent decision-making: for primates correct decisions are on average faster than errors, whereas for rodents correct decisions are on average slower than errors. Both observations violate the predictions of the standard drift diffusion model. In one study this species difference was seen even when sensory task was matched such that rats were just as accurate and just as fast as humans in the task, and even among subjects with low bias or lapse and comparable accuracy and speed (Shevinsky and Reinagel 2019).

We do not presume that the difference in response time of correct vs. error trials is functionally significant for either species; the difference is small and accounts for a small fraction of the variance in response time. The reason this effect is interesting is because it is places constraints on the underlying decision-making algorithms, and in particular, because it is inconsistent with DDM in its basic form.

Decreasing accuracy with response time has been widely reported in both humans and nonhuman primates (Roitman and Shadlen 2002, Palmer, Huk et al. 2005) and has been explained by a number of competing models (Ditterich 2006, Ditterich 2006, Ratcliff and McKoon 2008, Rao 2010, Hanks, Mazurek et al. 2011, Huang and Rao 2013, Ratcliff and Starns 2013, Tajima, Drugowitsch et al. 2016). It was only recently appreciated that accuracy increases with response time in this type of task in rats (Reinagel 2013, Reinagel 2013, Shevinsky and Reinagel 2019, Sriram, Li et al. 2020), and it remains unclear which of those models can accommodate this observation as well. In this study we showed that either parameter noise (Ratcliff and Tuerlinckx 2002, Ratcliff and McKoon 2008) or systematic parameter biases could explain the observed interaction between response time and accuracy in either species. Similar effects might be found in other related decision-making models.

On the models explored here, greater variability in the starting point of evidence accumulation would produce the effect seen in rats, whereas greater variability in the drift rate of evidence accumulation would produce the effect seen in humans. We do not know why rodents and primates should differ in this way. It could be, for example, that drift rate is modulated by top-down effects arising in cortex, while starting point is modulated by bottom-up effects arising subcortically, and species differ in the relative strength of these influences. Or perhaps some kinds of bias act on starting point while others act on drift rate, and species differ in which kinds of bias are stronger.

### Can context account for variability?

Although stochastic trial-by-trial variability of parameters could explain the effects of interest (Fig 3), systematic variations can also do so. We demonstrate this for simple cases of response side bias or history-dependent bias (Fig 4). Response bias is more prevalent in rats than in humans, but correct trials have longer RT than errors even in rats with no bias (Shevinsky and Reinagel 2019). In any case, these simulations show that response side bias would only produce the rat-like pattern if that bias impacted starting point to a greater degree than drift rate.

It is known that decisions in this type of task can be biased by the previous trial’s stimulus, response, and outcome in mice (Busse, Ayaz et al. 2011, Hwang, Dahlen et al. 2017, Roy, Bak et al. 2021), rats (Lavan, McDonald et al. 2011, Roy, Bak et al. 2021), non-human primates (Sugrue, Corrado et al. 2004), and humans (Goldfarb, Wong-Lin et al. 2012, Roy, Bak et al. 2021), reviewed in (Frund, Wichmann et al. 2014). Such history-dependent biases can be strong without causing an average side preference or an observable lapse rate (errors on strong stimuli). Species differ in the strength of such biases (Roy, Bak et al. 2021), but a difference in strength of bias does not determine whetherthe effect will be to make error trials earlier or later (Fig4L vs. 4P). This requires a difference in the computational site of action of bias.

In support of this idea, recent studies have traced variability to bias and history-dependent effects in both rodents and primates. In a go-nogo task, choice bias (conservative vs. liberal) in both mice and humans could be explained by bias in drift rate (de Gee, Tsetsos et al. 2020). In another study, choice history bias (repeat vs. alternate) was specifically linked to drift rate variability in humans (Urai, de Gee et al. 2019).

Fluctuations in arousal, motivation, satiety or fatigue could conceivably modulate decision thresholds or drift rates from trial to trial independently of either response side or trial history. (Note that in the model of Fig 2–3, fluctuations in the threshold separation parameter *a* are referred to the starting-point variability parameter *σ_z_* (Ratcliff and Tuerlinckx 2002, Ratcliff and McKoon 2008)). Such sources of variation may or may not be correlated with other measurable states, such as alacrity (e.g., latency to trial initiation, or in rodents the number or frequency of request licks), arousal (e.g., assessed by pupillometry), fatigue (the number of trials recently completed), satiety (amount of reward recently consumed), or frustration/success (fraction of recent trials penalized/rewarded). As models continue to include more of these effects, it will be of interest to determine how much of the observed behavioral variability is reducable to such deterministic components in each species, and whether those effects can be attributed differentially to starting point vs. drift rate effects in either decision-making models or neural recordings.

### Is parameter variability a bug or a feature?

To the extent that parameter variability is attributable to systematic influences rather than noise, a separate question would be whether this variability is adaptive or dysfunctional, in either species. It is possible that non-sensory influences shift the decision-making computation from trial to trial in a systematic and reproducible fashion that would be functionally adaptive in the context of natural behavior, even though we have artificially broken natural spatial and temporal correlations to render it maladaptive in our laboratory task.

For example, in nature some locations may be intrinsically more reward-rich, or very recent reward yields may be informative about the expected rewards at each location. In the real world, recently experienced visual motion might be highly predictive of the direction of subsequent motion stimuli. Therefore biasing either starting point or drift rate according to location or recent stimulus or reward history may be adaptive strategies under ecological constraints, for either or both species.

Consistent with this suggestion, decision biases of mice have been modeled as learned, continuously updated decision policies (Ashwood, Roy et al. 2020). Although the policy updates did not optimize expected reward in that study, the observed updates might still reflect hard-wired learning rules that would be optimal on average in natural contexts.

### Conclusions

It has been argued that neural computations underlying sensory decisions could integrate comparative information about incoming sensory stimuli (e.g., left vs. right motion signals), internal representations of prior probability (frequency of left vs. right motion trials) and the expected values (rewards or costs) associated with correct vs. incorrect decisions, in a common currency (Gold and Shadlen 2001, Gold and Shadlen 2007). On the premise that basic mechanisms of perceptual decision-making are likely to be conserved(Cesario, Johnson et al. 2020), fitting a single model to data from multiple species – especially where they differ in behavior – is a powerful way to develop and distinguish among alternative computational models (Urai, de Gee et al. 2019), and enables direct comparison of species.

## Methods

The data and code required to replicate all results in this manuscript are archived in a verified replication capsule (Reinagel and Nguyen 2021).

### Experimental data

No experiments were reported in this manuscript. Example human and rat data were previously published (Shevinsky and Reinagel 2019, Reinagel and Shevinsky 2020). Specifically, Fig 1F used rat fixed-coherence epoch 256 of that repository; Fig 1G-I used rat psychometric epoch 146. Fig 1J used human fixed-coherence epoch 63, and Fig1K-M used human psychometric epoch 81 from that data set. Fig 3D-M used the “AllEpochs” datasets.

To summarize the experiment briefly: the task was random dot coherent motion discrimination. When subjects initiated a trial, white dots appeared at random locations on a black screen and commenced to move. Some fraction of the dots (“signal”) moved at the same speed toward the rewarded response side. The others (“noise”) moved at random velocities. Subjects could respond at any time by a left vs. right keypress (human) or lick (rat). Correct responses were rewarded with money or water; error responses were penalized by a brief time-out. Stimulus strength was varied by the motion coherence (fraction of dots that were signal dots). Other stimulus parameters (e.g. dot size, dot density, motion speed, contrast) were chosen for each species to ensure that accuracy ranged from chance (50%) to perfect (100%) and response times ranged from ~500-2500ms for typical subjects of the species.

### Computational methods

The drift diffusion process was simulated according to the equation *X*(*t*) = *X*(*t*-1) ± δ with probability *p* of increasing and (1-*p*) of decreasing. Here *t* is the time point of the process, with time step τ in seconds; 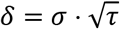 denotes the step size, where σ is the standard deviation of the Gaussian white noise of the diffusion; 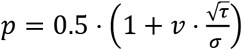, where *v* is the drift rate. The values for τ and σ were fixed at 0.1 msec and 1, respectively. For any trial, the process starts at a starting position *z*, sampled from a uniform distribution of range σ_z_, assumes a constant drift rate *v*, sampled from a normal distribution of standard deviation σ_v_, and continues until *X*(*t*) exceeds either threshold boundary. The non-decision-time *t*, sampled from a uniform distribution of range σ_t_, is added to the elapsed time to obtain the final RT associated with that trial.

We measured the interaction between accuracy and response time using the temporally local measure 〈RT_correct_ - RT_error_〉 introduced in (Shevinsky and Reinagel 2019). This method is preferred for real data because it is robust to non-trending non-stationarities that are commonly present in both human and rat data, not detected by traditional stationarity tests, and that could confound estimation of the effect of interest. The response time of each error trial is compared to a temporally adjacent correct trial of the same coherence, requiring a minimum distance of >3 trials to avoid sequential effects, and a maximum distance of 200 trials to avoid confounds due to long-range nonstationarity. For simulated data, where stationarity is guaranteed, the temporally local measure 〈RT_correct_ - RT_error_〉 and global measure 〈RT_correct_〉 - 〈RT_error_〉 are numerically equivalent.

We fit parameters of the model shown in Fig 2A to published human and rat datasets (Reinagel and Shevinsky 2020) using the Hierarchical Drift Diffusion Model (HDDM) package (Wiecki, Sofer et al. 2013). We emphasize that fitting the parameters of this model is problematic (Boehm, Annis et al. 2018). Our interpretation of the parameters (Fig 3D-M) is limited to asserting that these example parameters can produce human-like or rat-like effects, to the extent demonstrated. For further details of fitting, including scripts, raw output files and summary statistics of parameters, see Supplemental Materials.

## Supporting information

Supplemental Code: Model Fitting

