## Supplemental Code: Model Fitting for "Different forms of variability could explain a difference between human and rat decision making": README.txt

Supplementary material for Nguyen QN and Reinagel P (2022) Different Forms of Variability
Could Explain a Difference Between Human and Rat Decision Making. Front. Neurosci. 16:794681. doi: 10.3389/fnins.2022.794681

Model Fitting: Outputs

These are representative parameter fitting output files from HDDM after fitting the 
allEpochs datasets from:

Reinagel, Pamela; Shevinsky, Carly, 2020, "Human and Rat Motion Discrimination 
Reaction Time Task Data", https://doi.org/10.7910/DVN/ATMUIF, Harvard Dataverse, V1.

using the scripts provided in ../python scripts

The groups (humans, rats) were trained in separate runs so their parameters
are in different files. 

HDDM fitting was run for 102K iterations; statistics reported in these files 
are based on the final 50K, as indicated by the parameter 'length'

The columns in these files are:
parameter	if name does not contain "subj" it is a group posterior parameter
           if name contains "subj", the final numerical value is the epoch ID
mean: 		the mean over fitting iterations of the parameter value
std:		the std over fitting iterations of the parameter value
0.95 HDI lower:	the lower bound on the 95% High Density Interval for the parameter
0.95 HDI upper: 	the upper bound on the 95% High Density Interval for the parameter
*q:		the * percentile value of the parameter distribution (2.5, 25, 50, 75, 97.5)
mc err:		the Monte Carlo error (equivalent to standard error of the mean, for the mean in column 2)
length:		the number of iterations (from end of trace) included in calculation of these statistics
 
