## Supplemental Code: Model Fitting for "Different forms of variability could explain a difference between human and rat decision making": README.txt

Supplementary material for Nguyen QN and Reinagel P (2022) Different Forms of Variability
Could Explain a Difference Between Human and Rat Decision Making. Front. Neurosci. 16:794681. doi: 10.3389/fnins.2022.794681

The fitted parameters analyzed and plotted in Figure 3 of the manuscript are included in the CodeOcean repository: 
Pamela Reinagel, Quynh Nhu Nguyen (2022) Differential effects of variability could explain distinct human and rat deviations from DDM. Code Ocean 9600522.v2. doi: 10.24433/CO.9600522.v2.

Here we provide, in addition, methods used to fit the parameters using the HDDM package in Python.

The folder 'scripts' contains code and instructions to replicate the model fitting runs.  
The folder 'outputs' contains example model fitting run output files.

See the README file within each folder for more details.
