## Supplemental Code: Model Fitting for "Different forms of variability could explain a difference between human and rat decision making": analyze.py

```
import sys, os
from types import FunctionType

import numpy as np
from matplotlib.pylab import figure
import matplotlib.pyplot as plt

import pandas as pd
import pymc as pm
import pymc.progressbar as pbar
from . import utils

from .utils import interpolate_trace

from collections import OrderedDict

def plot_posterior_nodes(nodes, bins=50, lb=None, ub=None):
    """Plot interpolated posterior of a list of nodes.

    :Arguments:
        nodes : list of pymc.Node's
            List of pymc.Node's to plot the posterior of.
            These can be found in model.nodes_db.node.ix['param_name']
        bins : int (default=50)
            How many bins to use for computing the histogram.
        lb : float (default is to infer from data)
            Lower boundary to use for plotting.
        ub : float (default is to infer from data)
            Upper boundary to use for plotting.
    """
    figure()
    if lb is None:
        lb = min([min(node.trace()[:]) for node in nodes])
    if ub is None:
        ub = max([max(node.trace()[:]) for node in nodes])

    x_data = np.linspace(lb, ub, 300)

    for node in nodes:
        trace = node.trace()[:]
        #hist = interpolate_trace(x_data, trace, range=(trace.min(), trace.max()), bins=bins)
        hist = interpolate_trace(x_data, trace, range=(lb, ub), bins=bins)
        plt.plot(x_data, hist, label=node.__name__, lw=2.)

    leg = plt.legend(loc='best', fancybox=True)
    leg.get_frame().set_alpha(0.5)


def group_plot(model, params_to_plot=(), bins=50, samples=5000, save_to=None):
    def find_min_max(subj_block):
        # find global min and max for plotting
        min = np.inf
        max = -np.inf
        for name, subj in subj_block.iterrows():
            trace = subj['node'].trace()
            min = np.min([min, np.min(trace)])
            max = np.max([max, np.max(trace)])
        return min, max

    assert model.is_group_model, "group plot only works for group models."

    # select non-observed subject nodes
    subj_nodes = model.nodes_db[(model.nodes_db['observed'] == False) & (model.nodes_db['subj'] == True)]

    knode_names = subj_nodes.groupby(['knode_name', 'tag'])

    for (knode_name, tag), subj_block in knode_names:
        min, max = find_min_max(subj_block)

        # plot interpolated subject histograms
        #create figure
        print("plotting %s: %s" % (knode_name, tag))
        sys.stdout.flush()

        plt.figure()
        plt.title("%s: %s" % (knode_name, tag))
        x = np.linspace(min, max, 100)

        ############################################
        # plot subjects
        for name, subj_descr in subj_block.iterrows():
            trace = subj_descr['node'].trace()
            height = interpolate_trace(x, trace, range=(min, max), bins=bins)
            plt.plot(x, height, lw=1., label=str(np.int32(subj_descr['subj_idx'])))

        ###########################################
        # plot group distribution
        node = subj_descr['node']
        group_trace = np.empty(samples, dtype=np.float32)
        for sample in range(samples):
            # set parents to random value from their trace
            trace_pos = np.random.randint(0, len(node.trace()))
            for parent in node.extended_parents:
                parent.value = parent.trace()[trace_pos]
            group_trace[sample] = node.random()
            # TODO: What to do in case of deterministic (e.g. transform) node
            #except AttributeError:
            #    group_trace[sample] = node.parents.items()[0].random()

        height = interpolate_trace(x, group_trace, range=(min, max), bins=bins)
        plt.plot(x, height, '--', lw=2., label='group')

        ##########################################
        #legend and title
        leg = plt.legend(loc='best', fancybox=True)
        leg.get_frame().set_alpha(0.5)
        plt.gcf().canvas.set_window_title(knode_name)

        if save_to is not None:
            plt.savefig(os.path.join(save_to, "group_%s.png" % knode_name))
            plt.savefig(os.path.join(save_to, "group_%s.pdf" % knode_name))

def plot_all_pairwise(model):
    """Plot all pairwise posteriors to find correlations."""
    import scipy as sp
    from itertools import combinations
    #size = int(np.ceil(np.sqrt(len(data_deps))))
    fig = plt.figure()
    fig.subplots_adjust(wspace=0.4, hspace=0.4)
    # Loop through all pairwise combinations
    for i, (p0, p1) in enumerate(combinations(model.get_group_nodes()['node'], 2)):
        fig.add_subplot(6,6,i+1)
        plt.plot(p0.trace(), p1.trace(), '.')
        (a_s, b_s, r, tt, stderr) = sp.stats.linregress(p0.trace(), p1.trace())
        reg = sp.polyval((a_s, b_s), (np.min(p0.trace()), np.max(p0.trace())))
        plt.plot((np.min(p0.trace()), np.max(p0.trace())), reg, '-')
        plt.xlabel(p0.__name__)
        plt.ylabel(p1.__name__)

    plt.draw()

def gelman_rubin(models,num_samples,thin=5):
    """
    Calculate the gelman_rubin statistic (R_hat) for every stochastic in the model.
    (Gelman at al 2004, 11.4)
    Input:
        models - list of models
    """
    stochastics = models[0].get_stochastics()
    R_hat_dict = {}
##    num_samples = stochastics.node[0].trace().shape[0]
    num_chains = len(models)
    R_hat_dict['Number of samples'] = num_samples
    R_hat_dict['Number of chains'] = num_chains
    for name, stochastic in stochastics.iterrows():
        # Calculate mean for each chain
        samples = np.empty((num_chains, num_samples))
        for i,model in enumerate(models):
            samples[i,:] = model.nodes_db.ix[name, 'node'].trace(chain=None)[-num_samples::thin]

        R_hat_dict[name] = pm.diagnostics.gelman_rubin(samples)

    return R_hat_dict

R_hat = gelman_rubin

def check_geweke(model, assert_=True):
    # Test for convergence using geweke method
    for name, param in model.iter_stochastics():
        geweke = np.array(pm.geweke(param['node']))
        if np.any(np.abs(geweke[:,1]) > 2):
            msg = "Chain of %s not properly converged" % param
            if assert_:
                raise AssertionError(msg)
            else:
                print(msg)
            return False

    return True

def group_cond_diff(hm, node, cond1, cond2, threshold=0):
    """
    Compute the difference between different conditions in a group analysis.
    For each subject the function computes the difference between 'node' under
    condition 'cond1' to 'node' under condition 'cond2'.
    By assuming that each of the differences is normal distributed
    we can easily compute the group mean and group variance of the difference.
    Then the difference is compared to 'threshold' to compute the mass of the
    group pdf which is smaller than 'threshold'

    Input:
        hm - hierachical model
        node - name of node to be analyized
        cond1 - name of condition 1
        cond2 - name of condition 2
        threshold - see description

    Output:
        group_mean - group mean of the differnce
        group_var - group variance of the difference
        mass_under_threshold  - the mass of the group pdf which is smaller than threshold
    """
    import scipy as sp

    name = node
    node_dict = hm.params_include[name].subj_nodes
    n_subjs = hm._num_subjs

    #loop over subjs
    subj_diff_mean = np.zeros(n_subjs)
    subj_diff_std = np.zeros(n_subjs)
    for i_subj in range(n_subjs):
        #compute difference of traces
        name1 = node_dict[cond1][i_subj].__name__
        name2 = node_dict[cond2][i_subj].__name__
        trace1 = hm.mc.db.trace(name1)[:]
        trace2 = hm.mc.db.trace(name2)[:]
        diff_trace = trace1 - trace2

        #compute stats
        subj_diff_mean[i_subj] = np.mean(diff_trace)
        subj_diff_std[i_subj]= np.std(diff_trace)

    pooled_var = 1. / sum(1. / (subj_diff_std**2))
    pooled_mean = sum(subj_diff_mean / (subj_diff_std**2)) * pooled_var

    mass_under = sp.stats.norm.cdf(threshold,pooled_mean, np.sqrt(pooled_var))

    return pooled_mean, pooled_var, mass_under

def post_pred_compare_stats(sampled_stats, data_stats, evals=None):
    """Evaluate summary statistics of sampled sets.

    :Arguments:
        sampled_stats : dict
            Map of summary statistic names to distributions
        data_stats : dict
            Map of summary statistic names to the data distribution

    :Returns:
        pandas.DataFrame containing the eval results as columns.
    """

    from scipy.stats import scoreatpercentile, percentileofscore

    if evals is None:
        # Generate some default evals
        evals = OrderedDict()
        evals['observed'] = lambda x, y: y
        evals['mean'] = lambda x,y: np.mean(x)
        evals['std'] = lambda x,y: np.std(x)
        evals['SEM'] = lambda x, y: (np.mean(x) - y)**2
        evals['MSE'] = lambda x, y: np.mean((x - y)**2)
        evals['credible'] = lambda x, y: (scoreatpercentile(x, 97.5) > y) and (scoreatpercentile(x, 2.5) < y)
        evals['quantile'] = percentileofscore
        evals['mahalanobis'] = lambda x, y: np.abs(np.mean(x) - y)/np.std(x)
        #for q in [2.5, 25, 50, 75, 97.5]:
        #    key = str(q) + 'q'
        #    evals[key] = lambda x, y, q=q: scoreatpercentile(x, q)

    # Evaluate all eval-functions
    results = pd.DataFrame(index=list(sampled_stats.keys()), columns=list(evals.keys()) + ['NaN'],
                           dtype=np.float32)

    results.index.names = ['stat']
    for stat_name in sampled_stats:
        #update NaN column with the no. of NaNs and remove them
        s = sampled_stats[stat_name]
        results.ix[stat_name, 'NaN'] = sum(pd.isnull(s))
        s = s[np.isfinite(s)]
        if len(s) == 0:
            continue
        #evaluate
        for eval_name, func in evals.items():
            value = func(s, data_stats[stat_name])
            results.ix[stat_name, eval_name] = value

    return results.drop('NaN', axis=1)


def _post_pred_generate(bottom_node, samples=500, data=None, append_data=False):
    """Generate posterior predictive data from a single observed node."""
    datasets = []

    ##############################
    # Sample and generate stats
    for sample in range(samples):
        _parents_to_random_posterior_sample(bottom_node)
        # Generate data from bottom node
        sampled_data = bottom_node.random()
        if append_data and data is not None:
            sampled_data = sampled_data.join(data.reset_index(), lsuffix='_sampled')
        datasets.append(sampled_data)

    return datasets

def post_pred_gen(model, groupby=None, samples=500, append_data=False, progress_bar=True):
    """Run posterior predictive check on a model.

    :Arguments:
        model : kabuki.Hierarchical
            Kabuki model over which to compute the ppc on.

    :Optional:
        samples : int
            How many samples to generate for each node.
        groupby : list
            Alternative grouping of the data. If not supplied, uses splitting
            of the model (as provided by depends_on).
        append_data : bool (default=False)
            Whether to append the observed data of each node to the replicatons.
        progress_bar : bool (default=True)
            Display progress bar

    :Returns:
        Hierarchical pandas.DataFrame with multiple sampled RT data sets.
        1st level: wfpt node
        2nd level: posterior predictive sample
        3rd level: original data index

    :See also:
        post_pred_stats
    """
    results = {}

    # Progress bar
    if progress_bar:
        n_iter = len(model.get_observeds())
        bar = pbar.progress_bar(n_iter)
        bar_iter = 0
    else:
        print("Sampling...")

    if groupby is None:
        iter_data = ((name, model.data.ix[obs['node'].value.index]) for name, obs in model.iter_observeds())
    else:
        iter_data = model.data.groupby(groupby)

    for name, data in iter_data:
        node = model.get_data_nodes(data.index)

        if progress_bar:
            bar_iter += 1
            bar.update(bar_iter)

        if node is None or not hasattr(node, 'random'):
            continue # Skip

        ##############################
        # Sample and generate stats
        datasets = _post_pred_generate(node, samples=samples, data=data, append_data=append_data)
        results[name] = pd.concat(datasets, names=['sample'], keys=list(range(len(datasets))))

    if progress_bar:
        bar_iter += 1
        bar.update(bar_iter)

    return pd.concat(results, names=['node'])


def post_pred_stats(data, sim_datasets, stats=None, plot=False, bins=100, evals=None, call_compare=True):
    """Calculate a set of summary statistics over posterior predictives.

    :Arguments:
        data : pandas.Series

        sim_data : pandas.Series

    :Optional:
        bins : int
            How many bins to use for computing the histogram.
        stats : dict or function
            User-defined statistics to compute (by default mean and std are computed)
            and evaluate over the samples.
            :Example:
              * {'mean': np.mean, 'median': np.median}
              * lambda x: np.mean(x)
        evals : dict
            User-defined evaluations of the statistics (by default 95 percentile and SEM).
            :Example: {'percentile': scoreatpercentile}
        plot : bool
            Whether to plot the posterior predictive distributions.
        progress_bar : bool
            Display progress bar while sampling.
        field : string
            Which column name to run the stats on
        call_com,pare : bool (default=True)
            Whether to call post_pred_compare_stats. If False, return stats directly.
    """

    def _calc_stats(data, stats):
        out = {}
        for name, func in stats.items():
            out[name] = func(data)
        return out

    if stats is None:
        stats = OrderedDict((('mean', np.mean), ('std', np.std)))
    if isinstance(stats, FunctionType):
        stats = OrderedDict((('stat', stats),))

    data_stats = _calc_stats(data, stats)

    ###############################################
    # Initialize posterior sample stats container
    samples = len(sim_datasets)
    sampled_stats = {}
    sampled_stats = pd.DataFrame(index=sim_datasets.index.droplevel(2).unique(),
                                 columns=list(stats.keys()),
                                 dtype=np.float32)

    for i, sim_dataset in sim_datasets.groupby(level=(0, 1)):
        sampled_stat = _calc_stats(sim_dataset.values, stats)

        # Add it to the results container
        for name, value in sampled_stat.items():
            sampled_stats[name][i] = value

    if plot:
        from pymc.Matplot import gof_plot
        for name, value in sampled_stats.items():
            gof_plot(value, data_stats[name], bins=bins, name=name, verbose=0)

    if call_compare:
        return post_pred_compare_stats(sampled_stats, data_stats, evals=evals)
    else:
        return sampled_stats


def _parents_to_random_posterior_sample(bottom_node, pos=None):
    """Walks through parents and sets them to pos sample."""
    for i, parent in enumerate(bottom_node.extended_parents):
        if not isinstance(parent, pm.Node): # Skip non-stochastic nodes
            continue

        if pos is None:
            # Set to random posterior position
            pos = np.random.randint(0, len(parent.trace()))

        assert len(parent.trace()) >= pos, "pos larger than posterior sample size"
        parent.value = parent.trace()[pos]


def _plot_posterior_pdf_node(bottom_node, axis, value_range=None, samples=10, bins=100):
    """Calculate posterior predictive for a certain bottom node.

    :Arguments:
        bottom_node : pymc.stochastic
            Bottom node to compute posterior over.

        axis : matplotlib.axis
            Axis to plot into.

        value_range : numpy.ndarray
            Range over which to evaluate the likelihood.

    :Optional:
        samples : int (default=10)
            Number of posterior samples to use.

        bins : int (default=100)
            Number of bins to compute histogram over.

    """

    if value_range is None:
        # Infer from data by finding the min and max from the nodes
        raise NotImplementedError("value_range keyword argument must be supplied.")

    like = np.empty((samples, len(value_range)), dtype=np.float32)
    for sample in range(samples):
        _parents_to_random_posterior_sample(bottom_node)
        # Generate likelihood for parents parameters
        like[sample,:] = bottom_node.pdf(value_range)

    y = like.mean(axis=0)
    try:
        y_std = like.std(axis=0)
    except FloatingPointError:
        print("WARNING! %s threw FloatingPointError over std computation. Setting to 0 and continuing." % bottom_node.__name__)
        y_std = np.zeros_like(y)

    # Plot pp
    axis.plot(value_range, y, label='post pred', color='b')
    axis.fill_between(value_range, y-y_std, y+y_std, color='b', alpha=.8)

    # Plot data
    if len(bottom_node.value) != 0:
        axis.hist(bottom_node.value.values, normed=True, color='r',
                  range=(value_range[0], value_range[-1]), label='data',
                  bins=bins, histtype='step', lw=2.)

    axis.set_ylim(bottom=0) # Likelihood and histogram can only be positive

def plot_posterior_predictive(model, plot_func=None, required_method='pdf', columns=None, save=False, path=None,
                              figsize=(8,6), format='png', num_subjs=None, **kwargs):
    """Plot the posterior predictive distribution of a kabuki hierarchical model.

    :Arguments:

        model : kabuki.Hierarchical
            The (constructed and sampled) kabuki hierarchical model to
            create the posterior preditive from.

        value_range : numpy.ndarray
            Array to evaluate the likelihood over.

    :Optional:

        samples : int (default=10)
            How many posterior samples to generate the posterior predictive over.

        columns : int (default=3)
            How many columns to use for plotting the subjects.

        bins : int (default=100)
            How many bins to compute the data histogram over.

        figsize : (int, int) (default=(8, 6))

        save : bool (default=False)
            Whether to save the figure to a file.

        path : str (default=None)
            Save figure into directory prefix

        format : str or list of strings
            Save figure to a image file of type 'format'. If more then one format is
            givven them multiple files are created

        plot_func : function (default=_plot_posterior_pdf_node)
            Plotting function to use for each observed node
            (see default function for an example).

    :Note:

        This function changes the current value and logp of the nodes.

    """

    if plot_func is None:
        plot_func = _plot_posterior_pdf_node

    observeds = model.get_observeds()

    if columns is None:
        # If there are less than 3 items to plot per figure,
        # only use as many columns as there are items.
        max_items = max([len(i[1]) for i in
                         observeds.groupby('tag').groups.items()])
        columns = min(3, max_items)

    # Plot different conditions (new figure for each)
    for tag, nodes in observeds.groupby('tag'):
        fig = plt.figure(figsize=figsize)
        fig.suptitle(utils.pretty_tag(tag), fontsize=12)
        fig.subplots_adjust(top=0.9, hspace=.4, wspace=.3)

        # Plot individual subjects (if present)
        i = 0
        for subj_i, (node_name, bottom_node) in enumerate(nodes.iterrows()):
            i += 1
            if not hasattr(bottom_node['node'], required_method):
                continue # skip nodes that do not define the required_method

            ax = fig.add_subplot(np.ceil(len(nodes)/columns), columns, subj_i+1)
            if 'subj_idx' in bottom_node:
                ax.set_title(str(bottom_node['subj_idx']))

            plot_func(bottom_node['node'], ax, **kwargs)

            if num_subjs is not None and num_subjs >= i:
                break

        # Save figure if necessary
        if save:
            fname = 'ppq_' + '.'.join(str(tag_i) for tag_i in tag)
            if path is None:
                path = '.'
            if isinstance(format, str):
                format = [format]
            [fig.savefig('%s.%s' % (os.path.join(path, fname), x), format=x) for x in format]

        plt.close(fig)    

def geweke_problems(model, fname=None, **kwargs):
    """
    return a list of nodes who were detected as problemtic according to the geweke test
    Input:
        fname : string (deafult - None)
            Save result to file named fname
        kwargs : keywords argument passed to the geweke function
    """

    #search for geweke problems
    g = pm.geweke(model.mc)
    problems = []
    for node, output in g.items():
        values = np.array(output)[:,1]
        if np.any(np.abs(values) > 2):
            problems.append(node)

    #write results to file if needed
    if fname is not None:
        with open(fname, 'w') as f:
            for node in problems:
                f.write(node)

    return problems
```
