## Supplemental Code: Model Fitting for "Different forms of variability could explain a difference between human and rat decision making": hierarchical.py

```
#!/usr/bin/python

from copy import copy
import pickle
import sys

import numpy as np
from scipy.optimize import minimize, basinhopping

from collections import OrderedDict, defaultdict

import pandas as pd
import pymc as pm
import warnings
import matplotlib.pyplot as plt

from kabuki.utils import flatten
from . import analyze

class LnProb(object):
    def __init__(self, model):
        self.model = model

    def lnprob(self, vals): # vals is a vector of parameter values to try
        # Set each random variable of the pymc model to the value
        # suggested by emcee
        try:
            for val, (name, stoch) in zip(vals, self.model.iter_stochastics()):
                stoch['node'].set_value(val)
            logp = self.model.mc.logp
            return logp
        except pm.ZeroProbability:
            return -np.inf

    def __call__(self, *args, **kwargs):
        return self.lnprob(*args, **kwargs)

class Knode(object):
    def __init__(self, pymc_node, name, depends=(), col_name='',
                 subj=False, hidden=False, pass_dataframe=True, **kwargs):
        self.pymc_node = pymc_node
        self.name = name
        self.kwargs = kwargs
        self.subj = subj
        if isinstance(col_name, str):
            col_name = [col_name]

        self.col_name = col_name
        self.nodes = OrderedDict()
        self.hidden = hidden

        self.pass_dataframe = pass_dataframe

        #create self.parents
        self.parents = {}
        for (name, value) in self.kwargs.items():
            if isinstance(value, Knode):
                self.parents[name] = value

        # Create depends set and update based on parents' depends
        depends = set(depends)
        if self.subj:
            depends.add('subj_idx')
        depends.update(self.get_parent_depends())

        self.depends = sorted(list(depends))

        self.observed = 'observed' in kwargs

    def __repr__(self):
        return self.name

    def set_data(self, data):
        self.data = data

    def get_parent_depends(self):
        """returns the depends of the parents"""
        union_parent_depends = set()
        for name, parent in self.parents.items():
            union_parent_depends.update(set(parent.depends))
        return union_parent_depends

    def init_nodes_db(self):
        data_col_names = list(self.data.columns)
        node_descriptors = ['knode_name', 'stochastic', 'observed', 'subj', 'node', 'tag', 'depends', 'hidden']
        stats = ['mean', 'std', '2.5q', '25q', '50q', '75q', '97.5q', 'mc err']

        columns = node_descriptors + data_col_names + stats

        # create central dataframe
        self.nodes_db = pd.DataFrame(columns=columns)

    def append_node_to_db(self, node, uniq_elem):
        #create db entry for knode
        row = {}
        row['knode_name'] = self.name
        row['observed'] = self.observed
        row['stochastic'] = isinstance(node, pm.Stochastic) and not self.observed
        row['subj'] = self.subj
        row['node'] = node
        row['tag'] = self.create_tag_and_subj_idx(self.depends, uniq_elem)[0]
        row['depends'] = self.depends
        row['hidden'] = self.hidden

        row = pd.DataFrame(data=[row], columns=self.nodes_db.columns, index=[node.__name__])

        for dep, elem in zip(self.depends, uniq_elem):
            row[dep] = elem

        self.nodes_db = self.nodes_db.append(row)

    def create(self):
        """create the pymc nodes"""

        self.init_nodes_db()

        #group data
        if len(self.depends) == 0:
            grouped = [((), self.data)]
        else:
            grouped = self.data.groupby(self.depends)

        #create all the pymc nodes
        for uniq_elem, grouped_data in grouped:

            if not isinstance(uniq_elem, tuple):
                uniq_elem = (uniq_elem,)

            # create new kwargs to pass to the new pymc node
            kwargs = self.kwargs.copy()

            # update kwarg with the right parent
            for name, parent in self.parents.items():
                kwargs[name] = parent.get_node(self.depends, uniq_elem)

            #get node name
            tag, subj_idx = self.create_tag_and_subj_idx(self.depends, uniq_elem)
            node_name = self.create_node_name(tag, subj_idx=subj_idx)

            #get value for observed node
            if self.observed:
                if self.pass_dataframe:
                    kwargs['value'] = grouped_data[self.col_name] #.to_records(index=False)
                else:
                    kwargs['value'] = grouped_data[self.col_name].values #.to_records(index=False)

            # Deterministic nodes require a parent argument that is a
            # dict mapping parent names to parent nodes. Knode wraps
            # this; so here we have to fish out the parent nodes from
            # kwargs, put them into a parent dict and put that back
            # into kwargs, which will make pm.Determinstic() get a
            # parent dict as an argument.
            if self.pymc_node is pm.Deterministic:
                parents_dict = {}
                for name, parent in self.parents.items():
                    parents_dict[name] = parent.get_node(self.depends, uniq_elem)
                    kwargs.pop(name)
                kwargs['parents'] = parents_dict

                if self.observed:
                    kwargs['parents']['value'] = kwargs['value']


            # Deterministic nodes require a doc kwarg, we don't really
            # need that so if its not supplied, just use the name
            if self.pymc_node is pm.Deterministic and 'doc' not in kwargs:
                kwargs['doc'] = node_name

            node = self.create_node(node_name, kwargs, grouped_data)

            if node is not None:
                self.nodes[uniq_elem] = node
                self.append_node_to_db(node, uniq_elem)

    def create_node(self, node_name, kwargs, data):
        #actually create the node
        return self.pymc_node(name=node_name, **kwargs)

    def create_tag_and_subj_idx(self, cols, uniq_elem):
        uniq_elem = pd.Series(uniq_elem, index=cols)

        if 'subj_idx' in cols:
            subj_idx = uniq_elem['subj_idx']
            tag = uniq_elem.drop(['subj_idx']).values
        else:
            tag = uniq_elem.values
            subj_idx = None

        return tuple(tag), subj_idx


    def create_node_name(self, tag, subj_idx=None):
        # construct string that will become the node name
        s = self.name
        if len(tag) > 0:
            elems_str = '.'.join([str(elem) for elem in tag])
            s += "({elems})".format(elems=elems_str)
        if subj_idx is not None:
            s += ".{subj_idx}".format(subj_idx=subj_idx)

        return s

    def get_node(self, cols, elems):
        """Return the node that depends on the same elements.

        Called by the child to receive specific parent node.

        :Arguments:
            col_to_elem : dict
                Maps column names to elements.
                e.g. {'col1': 'elem1', 'col2': 'elem2', 'col3': 'elem3'}
        """

        col_to_elem = {}
        for col, elem in zip(cols, elems):
            col_to_elem[col] = elem

        # Find the column names that overlap with the ones we have
        overlapping_cols = intersect(cols, self.depends)

        # Create new tag for the specific elements we are looking for (that overlap)
        deps_on_elems = tuple([col_to_elem[col] for col in overlapping_cols])

        return self.nodes[deps_on_elems]

def intersect(t1, t2):
    # Preserves order, unlike set.
    return tuple([i for i in t2 if i in t1])

def test_subset_tuple():
    assert intersect(('a', 'b' , 'c'), ('a',)) == ('a',)
    assert intersect(('a', 'b' , 'c'), ('a', 'b')) == ('a', 'b')
    assert intersect(('a', 'b' , 'c'), ('a', 'c')) == ('a', 'c')
    assert intersect(('a', 'b' , 'c'), ('b', 'c')) == ('b', 'c')
    assert intersect(('c', 'b', 'a'), ('b', 'c')) == ('b', 'c')


class Hierarchical(object):
    """Creation of hierarchical Bayesian models in which each subject
    has a set of parameters that are constrained by a group distribution.

    :Arguments:
        data : numpy.recarray
            Input data with a row for each trial.
            Must contain the following columns:
              * 'rt': Reaction time of trial in seconds.
              * 'response': Binary response (e.g. 0->error, 1->correct)
            May contain:
              * 'subj_idx': A unique ID (int) of the subject.
              * Other user-defined columns that can be used in depends_on
                keyword.

    :Optional:
        is_group_model : bool
            If True, this results in a hierarchical
            model with separate parameter distributions for each
            subject. The subject parameter distributions are
            themselves distributed according to a group parameter
            distribution.

        depends_on : dict
            Specifies which parameter depends on data
            of a column in data. For each unique element in that
            column, a separate set of parameter distributions will be
            created and applied. Multiple columns can be specified in
            a sequential container (e.g. list)

            :Example:

            >>> depends_on={'param1':['column1']}

            Suppose column1 has the elements 'element1' and
            'element2', then parameters 'param1('element1',)' and
            'param1('element2',)' will be created and the
            corresponding parameter distribution and data will be
            provided to the user-specified method get_liklihood().

        trace_subjs : bool
             Save trace for subjs (needed for many
             statistics so probably a good idea.)

        plot_var : bool
             Plot group variability parameters

        In addition, the variable self.params must be defined as a
        list of Paramater().

    """

    def __init__(self, data, is_group_model=None, depends_on=None, trace_subjs=True,
                 plot_subjs=False, plot_var=False, group_only_nodes=()):
        # Init
        self.plot_subjs = plot_subjs
        self.depends_on = depends_on
        self.mc = None
        self.data = pd.DataFrame(data)
        self.group_only_nodes = group_only_nodes

        if not depends_on:
            depends_on = {}
        else:
            assert isinstance(depends_on, dict), "depends_on must be a dictionary."
            # Support for supplying columns as a single string
            # -> transform to list
            for key in depends_on:
                if isinstance(depends_on[key], str):
                    depends_on[key] = [depends_on[key]]
            # Check if column names exist in data
            for depend_on in depends_on.values():
                for elem in depend_on:
                    if elem not in self.data.columns:
                        raise KeyError("Column named %s not found in data." % elem)


        self.depends = defaultdict(lambda: ())
        for key, value in depends_on.items():
            self.depends[key] = value


        # Determine if group model
        if is_group_model is None:
            if 'subj_idx' in self.data.columns:
                if len(np.unique(data['subj_idx'])) != 1:
                    self.is_group_model = True
                else:
                    self.is_group_model = False
            else:
                self.is_group_model = False

        else:
            if is_group_model:
                if 'subj_idx' not in data.columns:
                    raise ValueError("Group models require 'subj_idx' column in input data.")

            self.is_group_model = is_group_model

        # Should the model incorporate multiple subjects
        if self.is_group_model:
            self._subjs = np.unique(data['subj_idx'])
            self._num_subjs = self._subjs.shape[0]
        else:
            self._num_subjs = 1

        self.num_subjs = self._num_subjs
        self.sampled = False
        self.dbname = 'ram'
        self.db = None

        self._setup_model()

    def _setup_model(self):
        # create knodes (does not build according pymc nodes)
        self.knodes = self.create_knodes()

        #add data to knodes
        for knode in self.knodes:
            knode.set_data(self.data)

        # constructs pymc nodes etc and connects them appropriately
        self.create_model()

    def __getstate__(self):
        from copy import deepcopy
        d = copy(self.__dict__)
        d['nodes_db'] = deepcopy(d['nodes_db'].drop('node', axis=1))
        d['depends'] = dict(d['depends'])
        #d['model_type'] = self.__class__

        if self.sampled:
            d['db'] = self.mc.db.__name__

            dbname = d['mc'].db.__name__
            if (dbname == 'ram'):
                    raise ValueError("db is 'ram'. Saving a model requires a database on disk.")
            elif (dbname == 'pickle'):
                    d['dbname'] = d['mc'].db.filename
            elif (dbname == 'txt'):
                    d['dbname'] = d['mc'].db._directory
            else: # hdf5, sqlite
                    d['dbname'] = d['mc'].db.dbname

        del d['mc']
        del d['knodes']

        return d

    def __setstate__(self, d):
        self.__dict__.update(d)
        self._setup_model()
        self.create_model()

        # backwards compat
        if not hasattr(self, 'sampled'):
            self.sampled = True

        if self.sampled:
            self.load_db(d['dbname'], db=d['db'])
            self.gen_stats()
        else:
            self.mcmc()

    def save(self, fname):
        """Save model to file.
        :Arguments:
           fname : str
              filename to save to

        :Notes:
            * Load models using kabuki.utils.load(fname).
            * You have to save traces to db, not RAM.
            * Uses the pickle protocol internally.
        """
        pickle.dump(self, open(fname, 'wb'))

    def create_knodes(self):
        raise NotImplementedError("create_knodes has to be overwritten")

    def create_model(self, max_retries=8):
        """Set group level distributions. One distribution for each
        parameter.

        :Arguments:
            retry : int
                How often to retry when model creation
                failed (due to bad starting values).
        """

        def _create():
            for knode in self.knodes:
                knode.create()

        for tries in range(max_retries):
            try:
                _create()
            except (pm.ZeroProbability, ValueError):
                continue
            break
        else:
            print("After %f retries, still no good fit found." %(tries))
            _create()

        # create node container
        self.create_nodes_db()

        # Check whether all user specified column names (via depends_on) where used by the depends_on.
        assert set(flatten(list(self.depends.values()))).issubset(set(flatten(self.nodes_db.depends))), "One of the column names specified via depends_on was not picked up. Check whether you specified the correct parameter value."

    def create_nodes_db(self):
        self.nodes_db = pd.concat([knode.nodes_db for knode in self.knodes])

    def draw_from_prior(self, update=False):
        if not update:
            values = self.values

        non_zero = True
        while non_zero:
            try:
                self.mc.draw_from_prior()
                self.mc.logp
                draw = copy(self.values)
                non_zero = False
            except pm.ZeroProbability:
                non_zero = True

        if not update:
            # restore original values
            self.set_values(values)

        return draw

    def map(self, runs=2, warn_crit=5, method='fmin_powell', **kwargs):
        """
        Find MAP and set optimized values to nodes.

        :Arguments:
            runs : int
                How many runs to make with different starting values
            warn_crit: float
                How far must the two best fitting values be apart in order to print a warning message

        :Returns:
            pymc.MAP object of model.

        :Note:
            Forwards additional keyword arguments to pymc.MAP().

        """

        from operator import attrgetter

        # I.S: when using MAP with Hierarchical model the subjects nodes should be
        # integrated out before the computation of the MAP (see Pinheiro JC, Bates DM., 1995, 2000).
        # since we are not integrating we get a point estimation for each
        # subject which is not what we want.
        if self.is_group_model:
            raise NotImplementedError("""Sorry, This method is not yet implemented for group models.
            you might consider using the approximate_map method""")

        maps = []

        for i in range(runs):
            # (re)create nodes to get new initival values.
            #nodes are not created for the first iteration if they already exist
            self.mc = pm.MAP(self.nodes_db.node)
            if i != 0:
                self.draw_from_prior()

            self.mc.fit(method, **kwargs)
            print(self.mc.logp)
            maps.append(self.mc)

        self.mc = None

        # We want to use values of the best fitting model
        sorted_maps = sorted(maps, key=attrgetter('logp'))
        max_map = sorted_maps[-1]

        # If maximum logp values are not in the same range, there
        # could be a problem with the model.
        if runs >= 2:
            abs_err = np.abs(sorted_maps[-1].logp - sorted_maps[-2].logp)
            if abs_err > warn_crit:
                print("Warning! Two best fitting MAP estimates are %f apart. Consider using more runs to avoid local minima." % abs_err)

        # Set values of nodes
        for max_node in max_map.stochastics:
            self.nodes_db.node.ix[max_node.__name__].set_value(max_node.value)

        return max_map


    def mcmc(self, assign_step_methods=True, *args, **kwargs):
        """
        Returns pymc.MCMC object of model.

        Input:
            assign_step_metheds <bool> : assign the step methods in params to the nodes

            The rest of the arguments are forwards to pymc.MCMC
        """

        self.mc = pm.MCMC(self.nodes_db.node.values, *args, **kwargs)

        self.pre_sample()

        return self.mc

    def pre_sample(self):
        pass

    def sample_emcee(self, nwalkers=500, samples=10, dispersion=.1, burn=5, thin=1, stretch_width=2., anneal_stretch=True, pool=None):
        import emcee
        import pymc.progressbar as pbar

        # This is the likelihood function for emcee
        lnprob = LnProb(self)

        # init
        self.mcmc()

        # get current values
        stochs = self.get_stochastics()
        start = [node_descr['node'].value for name, node_descr in stochs.iterrows()]
        ndim = len(start)

        def init_from_priors():
            p0 = np.empty((nwalkers, ndim))
            i = 0
            while i != nwalkers:
                self.mc.draw_from_prior()
                try:
                    self.mc.logp
                    p0[i, :] = [node_descr['node'].value for name, node_descr in stochs.iterrows()]
                    i += 1
                except pm.ZeroProbability:
                    continue
            return p0

        if hasattr(self, 'emcee_dispersions'):
            scale = np.empty_like(start)
            for i, (name, node_descr) in enumerate(stochs.iterrows()):
                knode_name = node_descr['knode_name'].replace('_subj', '')
                scale[i] = self.emcee_dispersions.get(knode_name, 0.1)
        else:
            scale = 0.1

        p0 = np.random.randn(ndim * nwalkers).reshape((nwalkers, ndim)) * scale * dispersion + start
        #p0 = init_from_priors()

        # instantiate sampler passing in the pymc likelihood function
        sampler = emcee.EnsembleSampler(nwalkers, ndim, lnprob, a=stretch_width, pool=pool)

        bar = pbar.progress_bar(burn + samples)
        i = 0

        annealing = np.linspace(stretch_width, 2, burn)
        sys.stdout.flush()

        for pos, prob, state in sampler.sample(p0, iterations=burn):
            if anneal_stretch:
                sampler.a = annealing[i]
            i += 1
            bar.update(i)

        #print("\nMean acceptance fraction during burn-in: {}".format(np.mean(sampler.acceptance_fraction)))
        sampler.reset()

        # sample
        try:
            for p, lnprob, lnlike in sampler.sample(pos,
                                                    iterations=samples,
                                                    thin=thin):
                i += 1
                bar.update(i)
        except KeyboardInterrupt:
            pass
        finally:
            print(("\nMean acceptance fraction during sampling: {}".format(np.mean(sampler.acceptance_fraction))))
            # restore state
            for val, (name, node_descr) in zip(start, stochs.iterrows()):
                node_descr['node'].set_value(val)

            # Save samples back to pymc model
            self.mc.sample(1, progress_bar=False) # This call is to set up the chains
            for pos, (name, node) in enumerate(stochs.iterrows()):
                node['node'].trace._trace[0] = sampler.flatchain[:, pos]

            return sampler

    def sample(self, *args, **kwargs):
        """Sample from posterior.

        :Note:
            Forwards arguments to pymc.MCMC.sample().

        """

        # Fetch out arguments for db backend
        db = kwargs.pop('db', 'ram')
        dbname = kwargs.pop('dbname', None)

        # init mc if needed
        if self.mc == None:
            self.mcmc(db=db, dbname=dbname)

        # suppress annoying warnings
        if ('hdf5' in dir(pm.database)) and \
           isinstance(self.mc.db, pm.database.hdf5.Database):
            warnings.simplefilter('ignore', pm.database.hdf5.tables.NaturalNameWarning)

        # sample
        self.mc.sample(*args, **kwargs)

        self.sampled = True

        self.gen_stats()
        return self.mc

    @property
    def logp(self):
        if self.mc is None:
            raise AttributeError('self.mc not set. Call mcmc().')
        return self.mc.logp

    @property
    def dic_info(self):
        """returns information about the model DIC."""

        info = {}
        try:
            info['DIC'] = self.mc.DIC
            info['deviance'] = np.mean(self.mc.db.trace('deviance')(), axis=0)
            info['pD'] = info['DIC'] - info['deviance']
        except pm.ZeroProbability:
            info['DIC'] = np.nan
            info['deviance'] = np.nan
            info['pD'] = np.nan

        return info

    @property
    def dic(self):
        """Deviance Information Criterion.
        """
        return self.dic_info['DIC']

    @property
    def aic(self):
        """Akaike Information Criterion.
        """
        if self.is_group_model:
            raise NotImplementedError('AIC can only be computed for non-hierarchical models. See dic.')
        k = len(self.get_stochastics())
        logp = sum([x.logp for x in self.get_observeds()['node']])
        return 2 * k - 2 * logp

    @property
    def bic(self):
        """Bayesian Information Criterion.
        """
        if self.is_group_model:
            raise NotImplementedError('BIC can only be computed for non-hierarchical models. See dic.')
        k = len(self.get_stochastics())
        n = len(self.data)
        logp = sum([x.logp for x in self.get_observeds()['node']])
        return -2 * logp + k * np.log(n)

    def _output_stats(self, stats_str, fname=None):
        """
        used by print_stats and print_group_stats to print the stats to the screen
        or to file
        """
        info = self.dic_info
        if fname is None:
            print(stats_str)
            print("DIC: %f" % info['DIC'])
            print("deviance: %f" % info['deviance'])
            print("pD: %f" % info['pD'])
        else:
            with open(fname, 'w') as fd:
                fd.write(stats_str)
                fd.write("\nDIC: %f\n" % info['DIC'])
                fd.write("deviance: %f\n" % info['deviance'])
                fd.write("pD: %f" % info['pD'])


    def gen_stats(self, fname=None, print_hidden=False, start=0, **kwargs):
        """print statistics of all variables
        Input (optional)
            fname <string> - the output will be written to a file named fname
            print_hidden <bool>  - print statistics of hidden nodes
        """
        self.append_stats_to_nodes_db(start=start)

        sliced_db = self.nodes_db.copy()

        # only print stats of stochastic, non-observed nodes
        if not print_hidden:
            sliced_db = sliced_db[(sliced_db['observed'] == False) & (sliced_db['hidden'] == False)]
        else:
            sliced_db = sliced_db[(sliced_db['observed'] == False)]

        stat_cols  = ['mean', 'std', '0.95 HDI lower', '0.95 HDI upper', '2.5q', '25q', '50q', '75q', '97.5q', 'mc err', 'length']

        for node_property, value in kwargs.items():
            sliced_db = sliced_db[sliced_db[node_property] == value]

        sliced_db = sliced_db[stat_cols]

        return sliced_db

    def print_stats(self, fname=None, print_hidden=False, **kwargs):
        """print statistics of all variables
        Input (optional)
            fname <string> - the output will be written to a file named fname
            print_hidden <bool>  - print statistics of hidden nodes
        """

        sliced_db = self.gen_stats(fname=fname, print_hidden=print_hidden, **kwargs)
        self._output_stats(sliced_db.to_string(), fname)

    def append_stats_to_nodes_db(self, *args, **kwargs):
        """
        smart call of MCMC.stats() for the model
        """
        try:
            nchains = self.mc.db.chains
        except AttributeError:
            raise ValueError("No model found.")

        #check which chain is going to be "stat"
        if 'chain' in kwargs:
            i_chain = kwargs['chain']
        else:
            i_chain = nchains

        #update self._stats
        self._stats = self.mc.stats(*args, **kwargs)
        self._stats_chain = i_chain

        #add/overwrite stats to nodes_db
        for name, i_stats in self._stats.items():
            if self.nodes_db.loc[name, 'hidden']:
                continue
            self.nodes_db.loc[name, 'mean']             = i_stats['mean']
            self.nodes_db.loc[name, 'std']              = i_stats['standard deviation']
            self.nodes_db.loc[name, '0.95 HDI lower']   = i_stats['95% HPD interval'][0]
            self.nodes_db.loc[name, '0.95 HDI upper']   = i_stats['95% HPD interval'][1]
            self.nodes_db.loc[name, '2.5q']             = i_stats['quantiles'][2.5]
            self.nodes_db.loc[name, '25q']              = i_stats['quantiles'][25]
            self.nodes_db.loc[name, '50q']              = i_stats['quantiles'][50]
            self.nodes_db.loc[name, '75q']              = i_stats['quantiles'][75]
            self.nodes_db.loc[name, '97.5q']            = i_stats['quantiles'][97.5]
            self.nodes_db.loc[name, 'mc err']           = i_stats['mc error']
            self.nodes_db.loc[name, 'length']           = i_stats['n']


    def load_db(self, dbname, verbose=0, db='sqlite'):
        """Load samples from a database created by an earlier model
        run (e.g. by calling .mcmc(dbname='test'))

        :Arguments:
            dbname : str
                File name of database
            verbose : int <default=0>
                Verbosity level
            db : str <default='sqlite'>
                Which database backend to use, can be
                sqlite, pickle, hdf5, txt.
        """


        if db == 'sqlite':
            db_loader = pm.database.sqlite.load
        elif db == 'pickle':
            db_loader = pm.database.pickle.load
        elif db == 'hdf5':
            db_loader = pm.database.hdf5.load
        elif db == 'txt':
            db_loader = pm.database.txt.load

        # Ignore annoying sqlite warnings
        warnings.simplefilter('ignore', UserWarning)

        # Open database
        db = db_loader(dbname)

        # Create mcmc instance reading from the opened database
        self.mc = pm.MCMC(self.nodes_db.node, db=db, verbose=verbose)

        # Not sure if this does anything useful, but calling for good luck
        self.mc.restore_sampler_state()

        return self

    def plot_posteriors(self, params=None, plot_subjs=False, save=False, **kwargs):
        """
        plot the nodes posteriors
        Input:
            params (optional) - a list of parameters to plot.
            plot_subj (optional) - plot subjs nodes
            kwargs (optional) - optional keywords to pass to pm.Matplot.plot

        TODO: add attributes plot_subjs and plot_var to kabuki
        which will change the plot attribute in the relevant nodes
        """

        # should we save the figures
        kwargs.pop('last', None)

        if isinstance(params, str):
             params = [params]

        # loop over nodes and for each node if it
        for (name, node) in self.iter_non_observeds():
            if (params is None) or (node['knode_name'] in params): # plot params if its name was mentioned
                if not node['hidden']: # plot it if it is not hidden
                    plot_value = node['node'].plot
                    if (plot_subjs and node['subj']): # plot if it is a subj node and plot_subjs==True
                        node['node'].plot = True
                    if (params is not None) and  (node['knode_name'] in params): # plot if it was sepecficily mentioned
                        node['node'].plot = True
                    pm.Matplot.plot(node['node'], last=save, **kwargs)
                    node['node'].plot = plot_value
                    plt.close()

    def plot_posteriors_conditions(self, *args, **kwargs):
        """
        Plot all group posteriors listed in depends_on on individual graphs.

        Forwards arguments to kabuki.analyze.plot_posterior_nodes.
        """
        group_nodes = self.get_group_nodes()
        for dep in self.depends_on.keys():
            nodes = group_nodes.ix[group_nodes.knode_name == dep]
            if all(nodes.hidden == True):
                continue
            analyze.plot_posterior_nodes(nodes['node'], *args, **kwargs)

    def get_observeds(self):
        return self.nodes_db[self.nodes_db.observed == True]

    def iter_observeds(self):
        nodes = self.get_observeds()
        for node in nodes.iterrows():
            yield node

    def get_non_observeds(self):
        return self.nodes_db[self.nodes_db.observed == False]

    def iter_non_observeds(self):
        nodes = self.get_non_observeds()
        for node in nodes.iterrows():
            yield node

    def iter_stochastics(self):
        nodes = self.get_stochastics()
        for node in nodes.iterrows():
            yield node

    def get_stochastics(self):
        return self.nodes_db[self.nodes_db.stochastic == True]

    def get_subj_nodes(self, stochastic=True):
        select = (self.nodes_db['subj'] == True) & \
                 (self.nodes_db['stochastic'] == stochastic)

        return self.nodes_db[select]

    def iter_subj_nodes(self, **kwargs):
        nodes = self.get_subj_nodes(**kwargs)
        for node in nodes.iterrows():
            yield node

    def get_group_nodes(self, stochastic=True):
        select = (self.nodes_db['subj'] == False) & \
                 (self.nodes_db['stochastic'] == stochastic)

        return self.nodes_db[select]

    def iter_group_nodes(self, **kwargs):
        nodes = self.get_group_nodes(**kwargs)
        for node in nodes.iterrows():
            yield node

    def get_group_traces(self):
        """Returns a DataFrame containing traces of all stochastic
        group nodes in the model.
        """
        return pd.DataFrame({i.__name__: i.trace() for i in self.get_group_nodes().node})

    def get_traces(self):
        """Returns a DataFrame containing traces of all stochastic
        nodes in the model.

        :Note: It is quite easy to then save this trace to csv by
        calling model.get_traces().to_csv('samples.csv')
        """
        return pd.DataFrame({i.__name__: i.trace() for i in self.get_stochastics().node})

    def get_data_nodes(self, idx):
        data_nodes = []
        for name, node_descr in self.iter_observeds():
            node = node_descr['node']
            if set(idx).issubset(set(node.value.index)):
                data_nodes.append(node)

        if len(data_nodes) != 1:
            raise NotImplementedError("Supply a grouping so that at most 1 observed node codes for each group.")

        return data_nodes[0]

    def __getitem__(self, name):
        return self.nodes_db.ix[name]['node']

    @property
    def values(self):
        values = OrderedDict()
        for (name, node) in self.iter_non_observeds():
            if node['node'].value.shape == ():
                values[name] = node['node'].value[()]

        return values

    def set_values(self, new_values):
        """
        set values of nodes according to new_values
        Input:
            new_values <dict> - dictionary of the format {'node_name1': new_value1, ...}
        """
        for (name, value) in new_values.items():
            self.nodes_db.ix[name]['node'].set_value(value)

    def find_starting_values(self, *args, **kwargs):
        """Find good starting values for the different parameters by
        optimization.

        For more options see approximate_map and map. Arguments are forwarded.
        """
        if self.is_group_model:
            self.approximate_map(*args, **kwargs)
        else:
            self.map(*args, **kwargs)

    def _partial_optimize(self, optimize_nodes, evaluate_nodes, fall_to_simplex=True, minimizer='Powell', use_basin=False, debug=False, minimizer_kwargs=None, basin_kwargs=None):
        """Optimize part of the model.

        :Arguments:
            nodes : iterable
                list nodes to optimize.
        """
        if minimizer_kwargs is None:
            minimizer_kwargs = {}
        if basin_kwargs is None:
            basin_kwargs = {}

        non_observeds = [x for x in optimize_nodes if not x.observed]

        init_vals = [node.value for node in non_observeds]

        # define function to be optimized
        def opt(values):
            if debug: print(values)
            for value, node in zip(values, optimize_nodes):
                node.set_value(value)
            try:
                logp_optimize = [node.logp for node in optimize_nodes]
                logp_evaluate = [node.logp for node in evaluate_nodes]
                neglogp = -np.sum(logp_optimize) - np.sum(logp_evaluate)
                if debug: print(neglogp)
                return neglogp
            except pm.ZeroProbability:
                if debug: print('Outside support!')
                return np.inf

        # optimize
        if use_basin:
            try:
                minimizer_kwargs_passed = {'method': minimizer, 'options': minimizer_kwargs}
                basinhopping(opt, init_vals, minimizer_kwargs=minimizer_kwargs_passed, **basin_kwargs)
            except:
                if fall_to_simplex:
                    print("Warning: Powell optimization failed. Falling back to simplex.")
                    minimizer_kwargs_passed = {'method': minimizer, 'options': minimizer_kwargs}
                    basinhopping(opt, init_vals, minimizer_kwargs=minimizer_kwargs_passed, **basin_kwargs)
                else:
                    raise
        else:
            try:
                minimize(opt, init_vals, method=minimizer, options=minimizer_kwargs)
            except:
                if fall_to_simplex:
                    print("Warning: Powell optimization failed. Falling back to simplex.")
                    minimize(opt, init_vals, method='Nelder-Mead', options=minimizer_kwargs)
                else:
                    raise


    def _approximate_map_subj(self, minimizer='Powell', use_basin=False, fall_to_simplex=True, debug=False, minimizer_kwargs=None, basin_kwargs=None):
        # Optimize subj nodes
        for subj_idx in self.nodes_db.subj_idx.dropna().unique():
            stoch_nodes = self.nodes_db.ix[(self.nodes_db.subj_idx == subj_idx) & (self.nodes_db.stochastic == True)].node
            obs_nodes = self.nodes_db.ix[(self.nodes_db.subj_idx == subj_idx) & (self.nodes_db.observed == True)].node
            self._partial_optimize(stoch_nodes, obs_nodes, fall_to_simplex=fall_to_simplex, minimizer=minimizer, use_basin=use_basin, debug=debug, minimizer_kwargs=minimizer_kwargs, basin_kwargs=basin_kwargs)

    def approximate_map(self, individual_subjs=True, minimizer='Powell', use_basin=False, fall_to_simplex=True, cycles=1, debug=False, minimizer_kwargs=None, basin_kwargs=None):
        """Set model to its approximate MAP.

        :Arguments:
            individual_subjs : bool <default=True>
                Optimize each subject individually.
            minimizer : str <default='Powell'>
                Optimize using Powell. See numpy.optimize.minimize.
                Other choice might be 'Nelder-Mead'
            use_basin : bool <default=True>
                Use basin hopping optimization to avoid local minima.
            fall_to_simplex : bool <default=True>
                should map try using simplex algorithm if powell method failes
            cycles : int <default=1>
                How many times to optimize the model.
                Since lower level nodes depend on higher level nodes,
                they might be estimated differently in a second pass.
            minimizer_kwargs : dict <default={}>
                Keyword arguments passed to minimizer.
                See scipy.optimize.minimize for options.
            basin_kwargs : dict <default={}>
                Keyword arguments passed to basinhopping.
                See scipy.optimize.basinhopping for options.
            debug : bool <default=False>
                Whether to print current values and neg logp at each
                iteration.

        """
        ###############################
        # In order to find the MAP of a hierarchical model one needs
        # to integrate over the subj nodes. Since this is difficult we
        # optimize the generations iteratively on the generation below.

        # only need this to get at the generations
        # TODO: Find out how to get this from pymc.utils.find_generations()
        m = pm.MCMC(self.nodes_db.node)
        generations_unsorted = m.generations
        generations_unsorted.append(self.get_observeds().node)
        # Filter out empty generations
        generations_unsorted = [gen for gen in generations_unsorted if len(gen) != 0]
        # Sort generations according to order of nodes_db
        generations = []
        for gen in generations_unsorted:
            generations.append([row.node for name, row in self.nodes_db.iterrows()
                                if name in [node.__name__ for node in gen]])

        for cyc in range(cycles):
            for i in range(len(generations)-1, 0, -1):
                if self.is_group_model and individual_subjs and (i == len(generations) - 1):
                    self._approximate_map_subj(fall_to_simplex=fall_to_simplex, minimizer=minimizer, use_basin=use_basin, debug=debug, minimizer_kwargs=minimizer_kwargs, basin_kwargs=basin_kwargs)
                    continue
                # Optimize the generation at i-1 evaluated over the generation at i
                self._partial_optimize(generations[i-1], generations[i], fall_to_simplex=fall_to_simplex, minimizer=minimizer, use_basin=use_basin, debug=debug, minimizer_kwargs=minimizer_kwargs, basin_kwargs=basin_kwargs)

        #update map in nodes_db
        self.nodes_db['map'] = np.NaN
        for name, value in self.values.items():
            try:
                self.nodes_db.loc[name, 'map'] = value
            # Some values can be series which we'll just ignore
            except (AttributeError, ValueError):
                pass
```
