## Supplemental Code: Model Fitting for "Different forms of variability could explain a difference between human and rat decision making": README.txt

Supplementary material for Nguyen QN and Reinagel P (2022) Different Forms of Variability
Could Explain a Difference Between Human and Rat Decision Making. Front. Neurosci. 16:794681. doi: 10.3389/fnins.2022.794681

Parameter Fitting Code

The networks are trained using the HDDM package, which can be found here:

http://ski.clps.brown.edu/hddm_docs/
along with instructions for other packages that must be installed for it to run

You must also replace the following files in the directory
...\Anaconda3\pkgs\kabuki-0.6.1-py35_0\Lib\site-packages\kabuki\
with the edited versions provided here:
analyze.py 
hierarchical.py
utils.py
(description of what was edited is below)

These are the scripts for fitting the model:
VariableDDMRatData.py
VariableDDMHumanData.py

which operate on the data files:
allEpochsRAT200502.csv
allEpochsHUMAN200502.csv

The data files may be found at:
Reinagel, P., and Shevinsky, C. 2020, "Human and Rat Motion 
Discrimination Reaction Time Task Data", 
https://doi.org/10.7910/DVN/ATMUIF, Harvard Dataverse. 

These scripts will run as they are, if you put the data files in the
C:/tmp/ directory and have write access to that directory.

This has been tested in Anaconda2/3-4.0.0-Win and HDDM version 0.6.0

exampe output files are found in ../outputs
=========================================================================================== 

To run the Python scripts on specific cores in a Windows PC:

1- Open Command Line (e.g. by typing 'cmd' in the Search bar)
2- In the Command Prompt, type: 
	start /affinity N ipython -i
(N: the hex equivalent of the decimal 2^n number corresponding to core n, n=0,1,2,...; 
	can check in Task manager > Performance, right-click > Change graph to > Logical processors
-i: interactive mode, good to access the script variables after the script crashes/finishes running)
3- In the IPython prompt that pops up, type (for example)
	exec(open('C:/tmp/StandardDDMRatData.py').read(),globals()) 
 
=========================================================================================== 
Description of changes that were made to the HDDM files listed above:

File: analyze.py (***Line numbers after 143 may be offsetted by 2)
	Function: plot_posterior_predictive (line 486)
		Edits: 1) (modification; line 568) [...] '.'.join(str(tag_i) for tag_i in tag)
			  2) (insertion; at the end, same indent level as 'if save:') plt.close(fig)

	Function: gelman_rubin (line 133)
		Edits:	1) (modification; line 133) gelman_rubin(models, num_samples, thin=5):
			2) (deletion; line 142)
			3) (insertion; after line 143, highest indent level) 	R_hat_dict['Number of samples'] = num_samples
										R_hat_dict['Number of chains'] = num_chains 
			4) (modification; line 150) [...] trace(chain=None)[-num_samples::thin]
		(***deprecated change, irrelevant for the newest version of the running script)

File: hierarchical.py (***Line numbers after 786 may be offsetted by 3)
	Function: plot_posterior (line 833)
		Edit:	(insertion; at the end, lowest indent level) plt.close()
	Function: save (line 402)
		Edit: 	(modification; line 413) [...] open(fname, 'wb'))
		(may have already been changed with later versions of kabuki)
	Function: gen_stats (line 727)
		Edit:	1) (modification; line 727) [...] start=0, **kwargs):
			2) (modification; line 733) self.append_stats_to_nodes_db(start=start)
			3) (modification; line 743) [...] 'std', '0.95 HDI lower', '0.95 HDI upper', [...] 'length']
	Function: append_stats_to_nodes_db (line 762)
		Edit: 	(insertion; after line 786, same indent level as the surroundings)	self.nodes_db.loc[name, '0.95 HDI lower']   = i_stats['95% HPD interval'][0]
            											self.nodes_db.loc[name, '0.95 HDI upper']   = i_stats['95% HPD interval'][1]
            											self.nodes_db.loc[name, 'length']           = i_stats['n']
File: utils.py
	Function: load (line 18)
		Edit:	(modification; line 23) [...] open(fname, 'rb'))
		(may have already been changed with later versions of kabuki)

==============================================================================================================================
