## Supplemental Code: Model Fitting for "Different forms of variability could explain a difference between human and rat decision making": utils.py

```
import pickle
import sys
import string
import copy

import numpy as np
import pandas as pd
import pymc as pm
from functools import reduce

def flatten(l):
    return reduce(lambda x, y: list(x)+list(y), l)

def pretty_tag(tag):
    return tag[0] if len(tag) == 1 else ', '.join(str(tag))

def load(fname):
    """Load a hierarchical model saved to file via
    model.save(fname)

    """
    model = pickle.load(open(fname, 'rb'))
    return model

def get_traces(model):
    """Returns recarray of all traces in the model.

    :Arguments:
        model : kabuki.Hierarchical submodel or pymc.MCMC model

    :Returns:
        trace_array : recarray

    """
    if isinstance(model, pm.MCMC):
        m = model
    else:
        m = model.mc

    nodes = list(m.stochastics)

    names = [node.__name__ for node in nodes]
    dtype = [(name, np.float) for name in names]
    traces = np.empty(nodes[0].trace().shape[0], dtype=dtype)

    # Store traces in one array
    for name, node in zip(names, nodes):
        traces[name] = node.trace()[:]

    return traces

def logp_trace(model):
    """
    return a trace of logp for model
    """

    #init
    db = model.mc.db
    n_samples = db.trace('deviance').length()
    logp = np.empty(n_samples, np.double)

    #loop over all samples
    for i_sample in range(n_samples):
        #set the value of all stochastic to their 'i_sample' value
        for stochastic in model.mc.stochastics:
            try:
                value = db.trace(stochastic.__name__)[i_sample]
                stochastic.value = value

            except KeyError:
                print("No trace available for %s. " % stochastic.__name__)

        #get logp
        logp[i_sample] = model.mc.logp

    return logp


def interpolate_trace(x, trace, range=(-1,1), bins=100):
    """Interpolate distribution (from samples) at position x.

    :Arguments:
        x <float>: position at which to evalute posterior.
        trace <np.ndarray>: Trace containing samples from posterior.

    :Optional:
        range <tuple=(-1,1): Bounds of histogram (should be fairly
            close around region of interest).
        bins <int=100>: Bins of histogram (should depend on trace length).

    :Returns:
        float: Posterior density at x.
    """

    import scipy.interpolate

    x_histo = np.linspace(range[0], range[1], bins)
    histo = np.histogram(trace, bins=bins, range=range, density=True)[0]
    interp = scipy.interpolate.InterpolatedUnivariateSpline(x_histo, histo)(x)

    return interp

def save_csv(data, fname, *args, **kwargs):
    """Save record array to fname as csv.

    :Arguments:
        data <np.recarray>: Data array to output.
        fname <str>: File name.

    :Notes:
        Forwards call to pandas DataFrame.to_csv

    :SeeAlso: load_csv
    """
    pd.DataFrame(data).to_csv(fname, *args, **kwargs)


def load_csv(*args, **kwargs):
    """Load record array from csv.

    :Arguments:
        fname <str>: File name.
        See pandas.read_csv()

    :Optional:
        See pandas.read_csv()

    :Note:
        Forwards call to pandas.read_csv()

    :SeeAlso: save_csv, pandas.read_csv()
    """
    return pd.read_csv(*args, **kwargs)


def set_proposal_sd(mc, tau=.1):
    for var in mc.variables:
        if var.__name__.endswith('var'):
            # Change proposal SD
            mc.use_step_method(pm.Metropolis, var, proposal_sd = tau)

    return

def stochastic_from_dist(*args, **kwargs):
    return pm.stochastic_from_dist(*args, dtype=np.dtype('O'),
                                   mv=True, **kwargs)

def concat_models(models, concat_traces=True):
    """Concatenate traces of multiple identical models into a new
    model containing all traces of the individual models.

    """
    # copy first model
    target_model = copy.deepcopy(models[0])
    target_stochs = target_model.get_stochastics()
    # append traces
    for i, model in enumerate(models[1:]):
        stochs = model.get_stochastics()
        for node, target_node in zip(stochs.node, target_stochs.node):
            assert node.__name__ == target_node.__name__, "Node names do not match. You have to pass identical models."
            if concat_traces:
                target_node.trace._trace[0] = np.concatenate([target_node.trace[:], node.trace[:]])
            else:
                target_node.trace._trace[i+1] = node.trace[:]

    target_model.gen_stats()

    return target_model


###########################################################################
# The following code is directly copied from Twisted:
# http://twistedmatrix.com/trac/browser/tags/releases/twisted-11.1.0/twisted/python/reflect.py
# For the license see:
# http://twistedmatrix.com/trac/browser/trunk/LICENSE
###########################################################################

class _NoModuleFound(Exception):
    """
    No module was found because none exists.
    """


class InvalidName(ValueError):
    """
    The given name is not a dot-separated list of Python objects.
    """


class ModuleNotFound(InvalidName):
    """
    The module associated with the given name doesn't exist and it can't be
    imported.
    """


class ObjectNotFound(InvalidName):
    """
    The object associated with the given name doesn't exist and it can't be
    imported.
    """

def _importAndCheckStack(importName):
    """
    Import the given name as a module, then walk the stack to determine whether
    the failure was the module not existing, or some code in the module (for
    example a dependent import) failing.  This can be helpful to determine
    whether any actual application code was run.  For example, to distiguish
    administrative error (entering the wrong module name), from programmer
    error (writing buggy code in a module that fails to import).

    @raise Exception: if something bad happens.  This can be any type of
    exception, since nobody knows what loading some arbitrary code might do.

    @raise _NoModuleFound: if no module was found.
    """
    try:
        try:
            return __import__(importName)
        except ImportError:
            excType, excValue, excTraceback = sys.exc_info()
            while excTraceback:
                execName = excTraceback.tb_frame.f_globals["__name__"]
                if (execName is None or # python 2.4+, post-cleanup
                    execName == importName): # python 2.3, no cleanup
                    raise excType(excValue).with_traceback(excTraceback)
                excTraceback = excTraceback.tb_next
            raise _NoModuleFound()
    except:
        # Necessary for cleaning up modules in 2.3.
        sys.modules.pop(importName, None)
        raise

def find_object(name):
    """
    Retrieve a Python object by its fully qualified name from the global Python
    module namespace.  The first part of the name, that describes a module,
    will be discovered and imported.  Each subsequent part of the name is
    treated as the name of an attribute of the object specified by all of the
    name which came before it.  For example, the fully-qualified name of this
    object is 'twisted.python.reflect.namedAny'.

    @type name: L{str}
    @param name: The name of the object to return.

    @raise InvalidName: If the name is an empty string, starts or ends with
        a '.', or is otherwise syntactically incorrect.

    @raise ModuleNotFound: If the name is syntactically correct but the
        module it specifies cannot be imported because it does not appear to
        exist.

    @raise ObjectNotFound: If the name is syntactically correct, includes at
        least one '.', but the module it specifies cannot be imported because
        it does not appear to exist.

    @raise AttributeError: If an attribute of an object along the way cannot be
        accessed, or a module along the way is not found.

    @return: the Python object identified by 'name'.
    """

    if not name:
        raise InvalidName('Empty module name')

    names = name.split('.')

    # if the name starts or ends with a '.' or contains '..', the __import__
    # will raise an 'Empty module name' error. This will provide a better error
    # message.
    if '' in names:
        raise InvalidName(
            "name must be a string giving a '.'-separated list of Python "
            "identifiers, not %r" % (name,))

    topLevelPackage = None
    moduleNames = names[:]
    while not topLevelPackage:
        if moduleNames:
            trialname = '.'.join(moduleNames)
            try:
                topLevelPackage = _importAndCheckStack(trialname)
            except _NoModuleFound:
                moduleNames.pop()
        else:
            if len(names) == 1:
                raise ModuleNotFound("No module named %r" % (name,))
            else:
                raise ObjectNotFound('%r does not name an object' % (name,))

    obj = topLevelPackage
    for n in names[1:]:
        obj = getattr(obj, n)

    return obj

######################
# END OF COPIED CODE #
######################

def centered_half_cauchy_rand(S, size):
    """sample from a half Cauchy distribution with scale S"""
    return abs(S * np.tan(np.pi * pm.random_number(size) - np.pi/2.0))

def centered_half_cauchy_logp(x, S):
    """logp of half Cauchy with scale S"""
    x = np.atleast_1d(x)
    if sum(x<0): return -np.inf
    return pm.flib.cauchy(x, 0, S) + len(x) * np.log(2)

HalfCauchy = pm.stochastic_from_dist(name="Half Cauchy",
                                     random=centered_half_cauchy_rand,
                                     logp=centered_half_cauchy_logp,
                                     dtype=np.double)

if __name__ == "__main__":
    import doctest
    doctest.testmod()
```
