## Supplemental Code: Model Fitting for "Different forms of variability could explain a difference between human and rat decision making": VariableDDMHumanData.py

```
# -*- coding: utf-8 -*-
"""
Created on Tue Jan  1 20:04:07 2019

This script will run HDDM model fit for 102K iterations,
updating intermediate results every 2K iterations.

Specifications (see Set Up below):
        INDIR: directory containing the input csv data file (default C:/tmp/)
        INFILE: csv file containing behavioral data in compatible format
	DATALABEL: concise string indicating input data file (affects output filename)
        OUTDIR: output directory (C:/tmp/, you must have write permission)
        RUNNUM: indexes repeat fitting runs (affects output filename)
        START: starting point of the sampling trace to include in summary statistics computation
        
Outputs:
        csv file containing variable DDM parameters
        pickle files containing the DDM object and the traces of its parameters
        csv file with DIC values of the model every 2K iterations
        png files: trace+histogram+autocorr plots of group level parameters
                   plots of simulated vs actual RT distributions (errors coded as negative RTs) 
          
Variable DDM Model:
        z=0
        includes sv/sz/st
        v depends on coherence
        uses informative priors (default in HDDM)

This code requires several edits to HDDM code first, see README.txt

@author: QN

Replication code for: Nguyen & Reinagel (2020)

"""

#================
# Set up
#================
INDIR       = 'C:/tmp/'
INFILE      = 'allEpochsHUMAN200502.csv'
DATALABEL   = 'human'
OUTDIR      = 'C:/tmp/'
RUNNUM      = '1'
START       = 1000      # affects trace summary statistics
thin        = 1         # affects trace plots
lenChunk    = 2000      # number of iterations between interim analysis
numChunks   = 50         # number of chunks, after first chunk


#================
# Instructions and code for installing HDDM can be obtained at
# http://ski.clps.brown.edu/hddm_docs/
#================
import os, sys, re, pickle
import matplotlib.pyplot as plt
import pymc as pm
import hddm


def fnProcessHDDMchunks(MODEL, RUNNUM, outFile, start=0, thin=1):
#================
# Helper function: constructs output file names, supports interim saving and plotting
#================
    # Compute mean, std, 95% HDI, and mcerr of the MCMC trace, starting from 'start'
    m_stats = MODEL.gen_stats(start=start) 
    m_stats.to_csv(outFile + '.csv')
    # Plot the RT distributions
    MODEL.plot_posterior_predictive(save=True, path='_'.join(['varDDM', RUNNUM, 'figures_RT']))
    plt.close("all")    # To save memory
    # More plots
    for (name,node) in MODEL.iter_non_observeds():
        if not node['hidden'] and not node['subj']:
            # Plot the full MCMC trace+histogram+autocorr
            pm.Matplot.plot(MODEL.mc.trace(name, chain=None)[:],
                            name, path='_'.join(['varDDM', RUNNUM, 'figures_tracesAllChains']))
            plt.close("all")
            # Plot the newest portion of the MCMC trace
            pm.Matplot.plot(node['node'].trace[::thin],
                            name, path='_'.join(['varDDM', RUNNUM, 'figures_tracesLastChain']))
            plt.close("all")

#================
# Main section
#================

print('\nFitting variable DDM to file {0:s}, attempt #{1:s}\n'.format(INFILE, RUNNUM))

# Read in the data file
inFilePath = os.path.join(INDIR, INFILE)
data = hddm.load_csv(inFilePath)
data = hddm.utils.flip_errors(data)
# Prepare the output folders and files
os.makedirs(OUTDIR, exist_ok=True)
os.chdir(OUTDIR)
outFile = '_'.join(['varDDM', DATALABEL, RUNNUM])         # output file stem
with open('LOGFILE.csv', 'a') as fid0:                  # append to the log file currently in OUTDIR
        fid0.write('{0:s},{1:s}\n'.format(
            inFilePath, outFile)) 
# Output image folders
p = re.sub(".csv", "", INFILE)                          # Clip the suffix in the file name
imRTPath        = os.path.join(p, '_'.join(['varDDM', RUNNUM, 'figures_RT']))
imTracePath1    = os.path.join(p, '_'.join(['varDDM', RUNNUM, 'figures_tracesAllChains']))
imTracePath2    = os.path.join(p, '_'.join(['varDDM', RUNNUM, 'figures_tracesLastChain']))
os.makedirs(imRTPath, exist_ok=True)
os.makedirs(imTracePath1, exist_ok=True)
os.makedirs(imTracePath2, exist_ok=True)
os.chdir(p)

#==============================================================
# HERE IS THE FUNCTION CALL DEFINING THE MODEL WE ARE FITTING:
m = hddm.HDDMTruncated(data,
                       include=('sv','sz','st'),
                       depends_on={'v':['species','coh'],
                                   'a':'species', 't':'species',
                                   'sv':'species', 'sz':'species', 'st':'species'})
m.find_starting_values()                                # OPTIONAL!!! May sometimes fail and crash the code
m.sample(lenChunk, dbname='_'.join([outFile, 'traces.db']), db='pickle')
#==============================================================

# Process the intermediate outputs
fnProcessHDDMchunks(m, RUNNUM, outFile, start=START, thin=thin)
totalIters = len(m.mc.trace('a_std', chain=None)[:])
with open('_'.join([outFile, 'dic.csv']), 'w') as fid:
    fid.write('DIC,totalIters\n')
    fid.write('{0:f},{1:d}\n'.format(m.dic, totalIters))

# Save the model
m.save(outFile)

# Continue fitting in chunks
for nIter in range(numChunks):
    print('\nCompleted {0:d} iterations\n'.format(totalIters))

    m.sample(lenChunk, dbname='_'.join([outFile, 'traces.db']), db='pickle')

    fnProcessHDDMchunks(m, RUNNUM, outFile, start=START, thin=thin)
    totalIters = len(m.mc.trace('a_std', chain=None)[:])
    with open('_'.join([outFile, 'dic.csv']), 'a') as fid:
        fid.write('{0:f},{1:d}\n'.format(m.dic, totalIters))

    m.save(outFile)
    
print('\nDone with file {0:s} after {1:d} iterations\n'.format(INFILE, totalIters))
```
