## Supplemental Materials for "Different forms of variability could explain a difference between human and rat decision making"

Supplementary material for: Nguyen QN and Reinagel P (2022) Different Forms of Variability Could Explain a Difference Between Human and Rat Decision Making. Front. Neurosci. 16:794681. doi: 10.3389/fnins.2022.794681

### *1. Supplementary statistics: Parameters fit to data (cf. Figure 3)*

| Dataset | Species | N | a | t | vmax | sz | sz/a | st | sv | sv/a |
|---|---|---|---|---|---|---|---|---|---|---|
| All | rat | 115 | 1.79+/-0.32 | 0.72+/-0.10 | 2.13+/-0.51 | 0.81+/-0.22 | 0.46+/-0.14 | 0.24+/-0.09 | 0.93+/-1.13 | 0.53+/-0.62 |
| All | human | 102 | 2.34+/-0.60 | 0.48+/-0.07 | 10.13+/-1.85 | 0.68+/-0.18 | 0.31+/-0.12 | 0.12+/-0.10 | 3.39+/-1.30 | 1.58+/-0.80 |
| Unbiased | rat | 11 | 1.54+/-0.28 | 0.67+/-0.10 | 1.91+/-0.47 | 0.63+/-0.24 | 0.42+/-0.16 | 0.24+/-0.10 | 0.47+/-0.46 | 0.35+/-0.43 |
| Unbiased | human | 51 | 2.27+/-0.56 | 0.48+/-0.07 | 10.68+/-2.10 | 0.72+/-0.18 | 0.34+/-0.13 | 0.11+/-0.08 | 3.64+/-1.33 | 1.74+/-0.88 |

**Table S1. Summary of fit parameters for the full datasets and the unbiased example subsets.**

All: fits to the “allEpochs” datasets of (Reinagel and Shevinsky 2020)
Unbiased: a subset of experiments identified from the “BestUnbiasedPsychometric” datasets
N: Number of experiments
a: threshold separation parameter (cf. Fig3 D,E)
t: non-decision time parameter (not plotted)
vmax: the drift rate of highest coherence (≥0.9 for rats, ≥0.24 for humans; not plotted)
sz: starting point variability parameter (not plotted)
sz/a: starting point variability as a fraction of threshold separation (cf. Fig3 F,G)
st: non-decision time variability parameter (not plotted)
sv: drift rate variability parameter (cf. Fig3 H,I)
sv/a: drift rate variability relative to threshold separation (not plotted)

The parameters reported are from the best (lowest deviance information criterion) of five independent fitting runs of the complete (“allEpochs”) dataset from each species. Summary statistics indicate mean±SD over experiments. For the unbiased subset, we report the parameters obtained for those experiments in the context of complete dataset fitting runs. The mean starting point $z$ was constrained to $z = 0$ in all cases. See section 2 below for fitting methods.

Note that most subjects contributed more than one experiment to the allEpochs datasets: the 115 rat experiments came from N=18 rats; the 102 human experiments came from N=63 subjects. In the Unbiased example set, each experiment is from a unique subject. Only 11/18 rat subjects and 51/63 human subjects are represented in the unbiased dataset because remaining subjects either had response bias, or had no psychometric experiments.

Experiments with too few error trials for the $\langle RT_c \rangle - \langle RT_e \rangle$ analysis are excluded from the upward bars of Fig 3 J-M, but the models fit to those experiments are included because more trials could be simulated to give the results in the lower bars.

In the source study, data collection was open-ended, and inclusion criteria for analysis were developed post-hoc; therefore, statistics intended for prospective hypothesis tests are inapplicable (Shevinsky and Reinagel 2019). Furthermore, we do not consider the fit parameters to be reliable measurements of real properties of the subjects (see section 3 below). Therefore, we think it would be overinterpreting the results to make any statistical claims about differences between the species on the basis of these parameter distributions.

### ***2. Supplementary methods: Model fitting** (cf. Figure 3)*

We fit the parameters of the variable-parameter model shown in Fig 2A (Ratcliff and McKoon 2008) to human and rat datasets (Reinagel and Shevinsky 2020) using HDDM (Wiecki, Sofer et al. 2013). The human and rat datasets were fit in separate runs. Thus, a value for each parameter was fit to each experiment in the dataset, mutually constrained by a species-specific group posterior. The Python scripts we used for fitting, and example training output files, are provided in Supplementary Code. Pertinent highlights are summarized here.

The models were trained for a pre-set 102,000 iterations with 2000 iteration burn-in. The adequacy of this training duration was assessed in preliminary runs by examining the asymptotic behavior of each parameter value and of the deviance information criterion (DIC) (Wiecki, Sofer et al. 2013). Some parameters were less confidently estimated – notably, the lower-coherence drift rates, the widths of the group posteriors, and the noise terms $\sigma_t$, $\sigma_z$, and $\sigma_v$ (Boehm, Annis et al. 2018). These parameter values often oscillated indefinitely with further training, and thus did not converge as judged by the highest density intervals (HDI) or the Gelman-Rubin $\hat{R}$ statistic (Gelman and Rubin 1992, Brooks and Gelman 1997). Pairs of parameters often fluctuated in a significantly correlated or anti-correlated way over iterations, indicating degeneracy of the solution. After 52,000 iterations, we are inclined to interpret any remaining fluctuations in the parameters as exploration of a flat region in the likelihood landscape (parameter combinations which produce about equally good fits).

The parameter values we report (Fig 3 D-I) and use for simulations (Fig 3 J-M) reflect the average value over the last 50,000 iterations of training. Despite the lack of convergence, these average values were well-estimated (Monte Carlo Error $MC_{err} < 2\%$ of parameter value) and reproducible (between-run coefficient of variation $CV < 2\%$ of parameter value). The total number of model parameters was 1485 (human) or 1217 (rats). The number of data points greatly exceeded the number of parameters: 102 trials/parameter (humans) or 225 trials/parameter (rats). In separate runs with split data, we verified that the parameter distributions and the prediction of the sign of $\langle RT_c - RT_e \rangle$ generalized upon cross-validation.

The purpose of fitting the model to datasets was to validate our conclusions from parameter sweeps. We did not pursue further optimizations for fitting this version of the model, because we don't favor this as the path forward for analyzing data. Rather, we think the best way to quantify and understand parameter variability in behavioral data will be to explicitly condition the parameters of DDM on measurable behaviors and states (cf. Fig 4), which is in the same spirit as other recent efforts (Urai, de Gee et al. 2019, Ashwood, Roy et al. 2020, Roy, Bak et al. 2021).

### ***3. Supplementary intuitions: Trial type pooling*** *(cf. Figure 4)*

Here we offer a different intuition for the effects of pooling left- and right-target trials. Analysis of **starting point bias** towards the right side are shown in Fig S1A. When we plot the dependence of mean response time on signed motion coherence, the curves for either leftward choices (magenta) or rightward choices (blue) are symmetric about coherence=0; within choices to either side, the RT is the same for correct responses (solid lines) and errors (dashed lines). Comparing the blue to magenta curves, however, we see that the blue curves always lie below the magenta curves, indicating that choices to the right (the preferred side) are faster than choices to the left, whether they are errors or correct responses. If we analyze $\langle RT_c \rangle - \langle RT_e \rangle$ for left-target and right-target trials separately (as in Fig 4D), the correct and error trials have unequal RTs in both cases (unlike a standard DDM). On the left side the errors are faster, whereas on the right side the correct trials are faster. To visualize this in Fig S1A, note that the separation between the blue and magenta curves is the same distance on both sides, but the curve for errors (dashed) lies below the curve for correct responses (solid) on the left, and above on the right. This explains why we obtain non-zero, symmetric values of $\langle RT_c \rangle - \langle RT_e \rangle$ on the two sides (Figure 4D).

When we pool the trials together, however, these effects do not cancel out because of asymmetries in the trial proportions. For example, at coherence=0.5, there are more fast trials (44+19=63% right-side choices, blue) than slow trials (31+6=37% left-side, magenta). The fast, preferred-side responses (blue) dominate both the errors (19/(19+6)=76%, dashed lines) and the correct responses

(44/(44+31)=59%, solid lines). But because the fast trial type dominates the error distribution more severely, the net effect is that errors are faster than correct trials on average (Fig 4E).

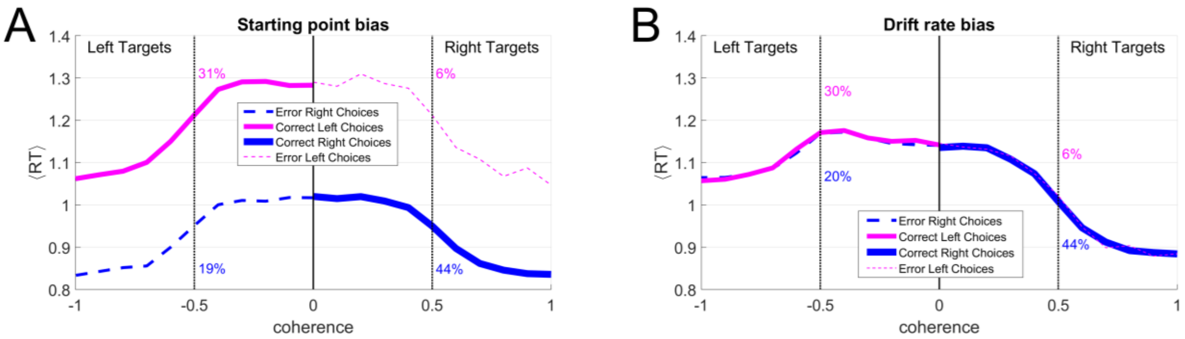


**Figure S1. Intuitions for the pooling effects in Figure 4.** Plots show the mean reaction time, <RT>, as a function of stimulus strength in simulated drift diffusion model runs. Negative coherence indicates leftward dot motion (left response target) and positive indicates rightward motion/target. The mean RT for each coherence is shown separately for four trial types: the left choices (magenta) and right choices (blue), errors (dashed lines) and correct responses (solid lines). The proportions of those four trial types are indicated by the thicknesses of the lines. Numbers indicate these proportions for an example case of motion coherence = 0.5 (black dotted lines). **(A)** Simulated rightward starting point bias. **(B)** Simulated rightward drift rate bias. See text for interpretation.

Results for a strong **drift rate bias** towards the right side are shown in Fig S1B. These curves are not symmetric left-to-right: left-motion trials are slower than right-motion trials of the same coherence. But the left-choice and right-choice curves overlap. Therefore, when we analyze either left-motion or right-motion trials separately, we find $\langle RT_c \rangle - \langle RT_e \rangle = 0$, as expected for a standard DDM (Fig 4I). When we combine trials of both motion directions in a pooled analysis, exactly half the trials are fast and half are slow (e.g. at coherence=0.5, 44+6=50% lower <RT>, 20+30=50% higher <RT>). But the error responses are dominated by slow, left-motion trials (20/(20+6)=77%, dashed lines), while the correct trials are dominated by fast, right-motion trials (44/(44+30)=59%, solid lines). This explains why on average, correct trials are faster: $\langle RT_c \rangle - \langle RT_e \rangle < 0$ (Fig 4J).

Note that for simplicity we have only discussed the mean RT in each condition. The full story is that the overlapping RT distributions of the four trial types are shifted early or late relative to one another in the direction summarized by the mean RT, such that all four trial types contribute unequally, in varying proportions depending on RT. The magnitudes of these effects change with coherence in both cases because the ratios of the four trial types change.

This same reasoning explains why in the general case, “fast errors” are produced by variability in the starting point, and “slow errors” by variability in the drift rate (Fig 3) as already pointed out by (Ratcliff and Rouder 1998). It also explains why any other factor causing the starting point to differ across trial types that are pooled in the analysis will produce a “fast error” phenotype (Fig 4N,O), while any other factor causing the drift rate to differ across pooled trial types will produce a “slow error” phenotype (Fig 4S,T).

### *4. Supplementary analysis (cf. Figure 4)*

For the benefit of those more accustomed to a different analysis, the same bias simulations analyzed in Figure 4 are re-analyzed an alternative way in Fig S2. In this analysis, trials are separated according to the animal’s behavioral choice (negative coherence indicating leftward choices) instead of according to the response target (direction of stimulus motion). The two analyses are equally informative (panels A-J), or equally uninformative (panels K-T), as the case may be. The main difference is that in this analysis, it is the starting point bias case that resolves to $\langle RT_c \rangle - \langle RT_e \rangle = 0$ in the side-conditioned analysis (Fig S2D, cf. Fig 4D) and the drift rate bias case that resolves to an equal but opposite non-zero difference (Fig S2I, cf. Fig 4I).

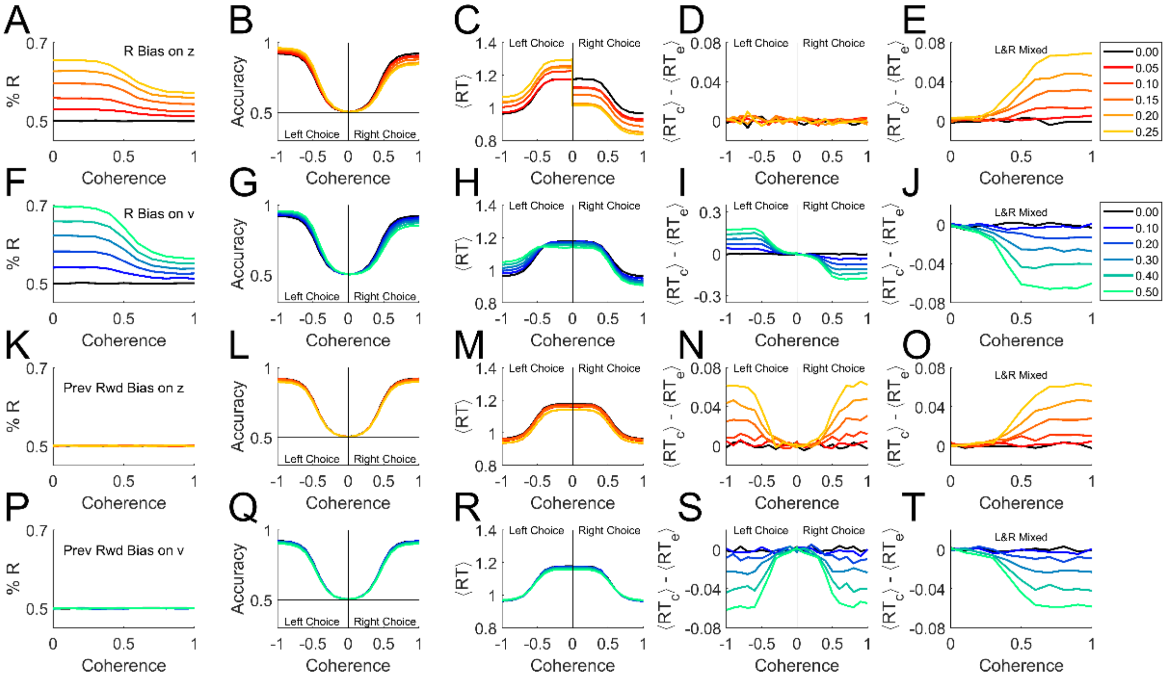


**Figure S2. Alternative analysis of simulated bias.** Simulations were performed with $\sigma_v = 0,\ \sigma_z = 0, \sigma_t = 0$ (i.e. the basic model in Fig 1) but with different forms of bias added. Trials were simulated with 50% Right targets (5x10$^5$ trials per coherence). **(A-E)** A Right-side bias was simulated by displacing the starting point $z$ toward the correct boundary on Right-target trials, or toward the error boundary for Left-target trials, by the amount indicated by color key at right. In contrast to Figure 4, in this case where coherence is signed, negative indicates leftward *behavioral choice* and positive indicates rightward behavioral choice. Within either behavioral choice direction, trials of both target locations (i.e. both motion directions) are pooled. If a coherence axis is unsigned, the left and right behavioral choices are pooled. **(A)** Percent Right responses, as a function of coherence (sensory stimulus strength), which determines the drift rate $v$. **(B)** Average accuracy of the response as a function of coherence. **(C)** Average response time as a function of coherence. **(D)** Difference between correct and error mean response times, $\langle RT_{correct} \rangle - \langle RT_{error} \rangle$, as a function of coherence. **(E)** Difference between correct and error mean responses times when left-choice and right-choice trials are pooled. **(F-J)** like A-E, but here bias was simulated by increasing the drift rate $v$ on R-target trials, or decreasing it on L-target trials, by the amount indicated in color key at right. **(K-O)** like A-E but here the starting point $z$ was displaced toward the side that was rewarded in previous trial (if any). Same color key as A-E. **(P-T)** like F-J but here the drift rate $v$ was increased if the target was on the side rewarded in previous trial, or decreased if the target was on the opposite side. Same color key as F-J.

### *References*

Ashwood, Z., N. Roy, J. H. Bak, T. I. B. Laboratory and J. W. Pillow (2020). Inferring learning rules from animal decision-making. Advances in Neural Information Processing Systems 33. H. Larochelle, M. Ranzato, R. Hadsell, M. F. Balcan and H. Lin, Curran Associates, Inc.: 3442-3453.
Boehm, U., J. Annis, M. J. Frank, G. E. Hawkins, A. Heathcote, D. Kellen, A. M. Krypotos, V. Lerche, G. D. Logan, T. J. Palmeri, D. van Ravenzwaaij, M. Servant, H. Singmann, J. J. Starns, A. Voss, T. V. Wiecki, D.

Matzke and E. J. Wagenmakers (2018). "Estimating across-trial variability parameters of the Diffusion Decision Model: Expert advice and recommendations." Journal of Mathematical Psychology **87**: 46-75.
Brooks, S. and A. Gelman (1997). "General methods for monitoring convergence of iterative simulations." J Computational and Graphical Statistics **7**: 434-455.
Gelman, A. and D. Rubin (1992). "Inference from iterative simulation using multiple sequences." Statistical Science **7**: 457-511.
Ratcliff, R. and G. McKoon (2008). "The diffusion decision model: theory and data for two-choice decision tasks." Neural Comput **20**(4): 873-922.
Ratcliff, R. and J. N. Rouder (1998). "Modeling response times for two-choice decisions." Psychological Science **9**(5): 347-356.
Reinagel, P. and C. A. Shevinsky (2020). Human and Rat Motion Discrimination Reaction Time Task Data. Harvard Dataverse.
Roy, N. A., J. H. Bak, L. International Brain, A. Akrami, C. D. Brody and J. W. Pillow (2021). "Extracting the dynamics of behavior in sensory decision-making experiments." Neuron **109**(4): 597-610 e596.
Shevinsky, C. A. and P. Reinagel (2019). "The Interaction Between Elapsed Time and Decision Accuracy Differs Between Humans and Rats." Frontiers in neuroscience **13**: 1211.
Urai, A. E., J. W. de Gee, K. Tsetsos and T. H. Donner (2019). "Choice history biases subsequent evidence accumulation." Elife **8**.
Wiecki, T. V., I. Sofer and M. J. Frank (2013). "HDDM: Hierarchical Bayesian estimation of the Drift-Diffusion Model in Python." Frontiers in Neuroinformatics **7**.
